# *bZIP63* misregulation affects growth and target gene expression under short-day photoperiods

**DOI:** 10.64898/2026.03.21.713353

**Authors:** Raphael de A. Campos, Pamela T. Carlson, Isis Sebastião, João G. P. Vieira, Cleverson C. Matiolli, Américo J. C. Viana, Michel Vincentz

## Abstract

Plant survival and growth depend partly on the ability to manage energy resources in response to changing environmental conditions. SnRK1 plays a central role in this process by restricting growth under energy-limiting conditions while promoting stress adaptation and survival. When activated, SnRK1 triggers transcriptional reprogramming that prioritizes energy-producing pathways. A key mediator of this response is the transcription factor bZIP63, whose activity is regulated by SnRK1-dependent phosphorylation. Given its roles in energy homeostasis and its interaction with the circadian clock, bZIP63 influences growth and is therefore a candidate component of the Metabolic Daylength Measurement (MDLM) system, which integrates starch and sucrose metabolism with circadian timing and photosynthetic duration to regulate vegetative growth under contrasting photoperiods. We show that 39 bZIP63 direct targets regulated by SnRK1 correspond to a subset of short-day–induced genes associated with the MDLM system and are downregulated in a *bZIP63* T-DNA mutant (*bzip63-2*) and/or in an RNAi-induced silencing line (*RNAiWs_L9*). Downregulation of these genes was more extensive in *RNAiWs_L9* than in *bzip63-2*, possibly due to the unexplained silencing of *BAM4*, a β-amylase that promotes starch degradation. Under short-day conditions, the frameshift mutant *bzip63-5* (Col-0), *bzip63-2* (Ws), and the *bzip1-1/bzip53-1/bzip63-5* (Col-0) triple mutant, which disrupts bZIP63 heterodimerization partners, showed similar deregulation of a subset of these genes and comparable growth inhibition, whereas both growth and gene deregulation were more strongly affected in *RNAiWs_L9*. We further show in two partially complemented *bzip63-2* lines that bZIP63 protein levels increase toward the end of the night and decline toward the end of the day, in synchrony with the diel oscillation of its transcript. Additional analyses of these lines, together with *bzip63-2* line overexpressing bZIP63, suggest that the timing and amplitude of bZIP63 accumulation contribute to shaping the expression profiles of a subset of the 39 MDLM-associated genes. Together, these findings indicate that bZIP63 participates in a regulatory network linking SnRK1 signaling, photoperiod-changes, and growth within the MDLM system.

## INTRODUCTION

Plants fine-tune growth and development by integrating external environmental cues, such as light, photoperiod, temperature, and stress, with internal signals, including circadian rhythms and energy status (Greenham and McClung, 2015; Baena-González and Hanson, 2017; Artins et al., 2024). Central to this integration are the Target of Rapamycin (TOR) and SNF1-related kinase 1 (SnRK1) evolutionary conserved kinases, which sense nutrient and energy availability to adjust growth demands (Robaglia et al., 2012; Baena-González and Hanson, 2017; Margalha et al., 2019; Artins et al., 2024). TOR signaling links metabolic and energy sufficiency to growth by promoting anabolic metabolism and meristem activity (Xiong et al., 2013; Dobrenel et al., 2016; Burkart and Brandizzi, 2021; Henriques et al., 2022). In contrast, SnRK1 is activated under energy-limiting and/or stress conditions, prioritizing energy conservation and survival by restraining growth in part by repressing TOR-dependent processes. (Baena-González et al., 2007; Crepin and Rolland, 2019; Peixoto and Baena-González, 2022; Baena-González and Lunn, 2020). Together, these two antagonistic pathways ensure appropriate growth and developmental responses to fluctuations in interconnected energy, metabolic, and stress conditions.

An important regulatory consequence of SnRK1 activation under energy deprivation is a transcriptional reprogramming that shifts metabolism from energy-consuming anabolic processes toward catabolic pathways. Catabolism enhances energy production by mobilizing alternative sources, including amino acids, lipids, and cellular components recycled through autophagy or alternative mitochondrial metabolic pathways (Baena-González et al., 2007; Pedrotti et al., 2018; Henninger et al., 2022). These transcriptional changes are mediated in part by the SnRK1α catalytic subunit, which phosphorylates the bZIP-type transcription factor *bZIP63* and consequently alters its heterodimerization preferences with group S1 bZIPs, thereby shaping starvation-induced gene expression programs (Mair et al. 2015; Pedrotti et al., 2018; Dröge Laser and Weiste, 2018; Van Leene et al., 2022). The SnRK1-bZIP63 regulatory module also acts in dark-induced stress/senescence (Mair et al., 2015; Pedrotti et al., 2018), contribute to early seedling establishment (Henninger et al., 2022), in lateral root priming after energy limitation (Muralidhara et al., 2021), in adjusting the circadian clock phase in response to sucrose by regulating the core clock component *PSEUDO-RESPONSE REGULATOR 7* (*PRR7*) in a *TREHALOSE 6-PHOSPHATE SYNTHASE1* (*TPS1*)-dependent manner (Frank et al., 2018). This later signaling pathway can partly explain *bZIP63*-mediated modulation of night starch decay to sustain growth. Moreover, the diel expression profile of *bZIP63* is dictated by both the circadian clock and cellular energy status, which in turn influences the expression of genes involved in starch degradation and adaptation to energy deprivation (Viana et al., 2021).

Starch metabolism and sucrose homeostasis have recently been implicated in a metabolic photoperiod measurement mechanism, termed MDLM system (Liu et al., 2021; Gendron and Staiger, 2023; Wang et al., 2024), which contributes to seasonal regulation of vegetative growth. The MDLM system integrates the circadian clock as a timer with the effective duration of photosynthesis and establishes photoperiod-specific gene expression programs characterized by defined oscillatory expression profile of multiple genes, including *PHLOEM PROTEIN2-A13* (*PP2-A13*) and *MYO-INOSITOL-1-PHOSPHATE SYNTHASE1* (*MIPS1*), which are essential for efficient vegetative growth under short- and long-day conditions, respectively (Liu et al., 2021; Wang et al., 2024).

Given its established roles in energy signaling, circadian regulation, and starch degradation, we hypothesized that *bZIP63* contributes to gene expression programs associated with the MDLM system. Here, we show that loss of bZIP63 activity disrupts the expression of a set of MDLM-associated genes and reduces growth more significantly under short-day conditions. Moreover, we show that diel oscillations in bZIP63 protein abundance closely match its mRNA dynamics, with a progressive increase during the night followed by decay during the daytime, suggesting that bZIP63 activity may be temporally fine-tuned in response to photoperiodic cues.

## MATERIAL AND METHODS

### Genotypes, growth conditions and sampling

The genotypes include *Arabidopsis thaliana* ecotypes Wassilewskija (Ws) and Columbia-0 (Col-0), the T-DNA insertion mutant *bzip63-2* (FLAG_610A08; Matiolli et al., 2011), and RNAi lines (*RNAiWs_L9*, *RNAiWs_L5*, and *RNAiWs_L7*) in which bZIP63 expression was silenced through an RNA interference approach (Frank et al., 2018). In addition, OX *bZIP63* lines (*bZIP63-ox3* and *bZIP63-ox4*) in which *bZIP63* is constitutively expressed, and the CO *bzip63-2* lines (*bZIP63-co1*, *bZIP63-co2* and *bZIP63-co3*) in which *bZIP63* expression is restored were analyzed.

Plants were grown in pots (8 cm diameter; 6 cm height; 301.59 cm³ volume) filled with a mixture of two parts of commercial substrate (Carolina Garden) to one part of vermiculite. After sowing, the pots were kept in darkness at 4 °C for 72 hours to break dormancy, and plants were grown at a constant temperature of approximately 20 °C with weekly watering. Light intensity was adjusted to 80 or 100 μmol m^−2^ s^−1^, and photoperiods were adjusted to short (8 hours of light and 16 hours of dark), equinoctial (12 hours of light and 12 hours of dark), or long (16 hours of light and 8 hours of dark). Every 15 days and 5 days before sampling a NPK nutrient solution was provided. The commercial NKP solution (Vithal for Plants and Balcony Flowers; density of 1.16 g L^-1^) consists of phosphoric acid, potassium nitrate, urea, and water, containing 8% total nitrogen, 4% phosphorus (P_2_O_5_) and 6% potassium (K_2_O) soluble in water. The solution was applied at final concentrations of 0.84 g L^-1^ nitrogen (total N), 0.42 g L^-1^ phosphate (P₂O₅), and 0.63 g L^-1^ potassium (K₂O).

Whole rosette leaves from plants grown 20 to 24 days after germination for short day photoperiod and 12 days for long day photoperiod were collected during the last 45 minutes of the night. Each sample consisted of rosettes from three to five individual plants grown in separate pots, with two to three replicates for protein analysis and three to five replicates for gene expression analysis per time point and genotype. Samples were immediately frozen in liquid nitrogen and subsequently stored at -80°C until nucleic acid extraction.

### DNA constructions and plant transformation

Construction to induce RNAi silencing of *bZIP63* were described in Frank et al., 2018.

bZIP63 cDNA fused to HA TAG in pENTR was transferred by Gateway cloning into the binary vector pH7FWG2 under the control of the CaMV 35S promoter and fused in C-terminal to Green Fluorescent Protein (GFP) to generate the 35S:HA:bZIP63:GFP construct in pH7FWG2 (Karimi et al., 2002). The 3341 bp *bZIP63* promoter sequence was obtained by amplification with primers P_bZIP63_Fw1 + P_SpeI_bZIP63_Rv1 (Table S3) and was digested with Hind-III and Spe-I resulting in a 2533 bp fragment. The CaMV 35S promoter was removed from 35S:HA:bZIP63:GFP by HindIII / Spe-I digestion and replaced by the 2533 bp *bZIP63* promoter fragment to generate 35S:HA:bZIP63:GFP within pH7FWG2. *bzip63-2* transformation was as described by Frank et al., 2018 and primary transformants were selected by hygromycin resistance. Homozygotic lines were selected for each construction based on hygromycin resistance and were genotyped.

### Generation of single, double, and triple mutants

Two *bZIP63* frame shift mutant lines in Col-0 ecotype, *bzip63-5* and *bzip63-15*, were generated using the CRISPR/Cas system by Vienna Bio Center Core Facilities (www.vbcf.ac.at). These mutations were confirmed through Sanger sequencing, showing insertions (indels) that caused frameshifts and premature stop codons (Figure S4). To generate double mutants *bzip1-1/bzip63-5* and *bzip53-1/bzip63-5*, crosses were made between *bzip63-5* and the T-DNA insertion mutants *bzip1-1* (SALK 059343; Dietrich et al., 2011; Figure S8) and *bzip53-1* (SALK_069883C; Dietrich et al., 2011; Figure S8). F1 and F2 plants were genotyped for *bzip1-1* and *bzip53-1* by PCR to detect T-DNA insertions, and for *bzip63-5* by sequencing. For the triple *bzip1-1/bzip53-1/bzip63-5* mutant, the double mutant *bzip1-1/bzip63-5* was crossed with *bzip53-1*. F1 and F2 plants were genotyped as described above.

### DNA Extraction, Conventional PCR, and Sequencing

Genomic DNA was extracted from leaves that had been previously macerated and stored at -80°C until processing. The extraction was carried out using the CTAB method with 100 mg of macerated material (Doyle & Doyle, 1990). DNA concentration was measured using a NanoVueTM spectrophotometer (GE Healthcare), and the integrity of the DNA was verified by electrophoresis on a 1% agarose gel.

Conventional PCR amplification of target regions was performed with specific primers and GoTaq® DNA Polymerase (Promega). PCR conditions were as follows: initial denaturation at 94 °C for 3 min, followed by 30 cycles of denaturation at 94 °C for 45 s, annealing at 55 °C for 30 s, and extension at 72 °C for 90 s, with a final extension at 72 °C for 10 min.

Sanger sequencing was carried out using the Genetic Analyzer 3500XL system (Life Technologies – Applied Biosystems). Sequencing reactions were performed using the BigDye™ Terminator v3.1 Cycle Sequencing Kit (Promega Biotech).

### RNA Extraction, DNase Treatment, and cDNA Synthesis

Leaves (∼100 mg) were macerated and used for total RNA extraction using the method described by Oñate-Sánchez and Vicente-Carbajosa (2008). RNA concentration was quantified with a NanoVueTM spectrophotometer (GE Healthcare), and RNA integrity was assessed by electrophoresis on a 1% (w/v) agarose gel with 5% formaldehyde, as described by Logemann et al. (1987).

DNAse treatment was performed using the TURBO DNA-free™ Kit (Invitrogen) and DNAse was inactivated with DNAse Inactivation Reagent as recommended by the manufacturer.

cDNA was synthesized using oligo-dT as a primer and ImProm-II™ Reverse Transcription Kit (Promega) for reverse transcription as recommended by the manufacturer.

### Quantitative Real-Time PCR (RT-qPCR)

The iTaq Universal SYBR Green Supermix Kit (Bio-Rad) was used for RT-qPCR of cDNA obtained from total RNA. Parameters for RT-qPCR included an initial denaturation at 95°C for 10 minutes, followed by 40 cycles of denaturation at 95°C for 15 seconds, and annealing and extension at 60°C for 1 minute, during which fluorescence readings were taken. Dissociation curves were obtained to check for non-specific amplifications and primer-dimer formation. Ct values were normalized to the reference gene *AT1G13320* or *AT3G18780* (Czechowski et al., 2005), and gene expression levels were calculated using the 2^-ΔΔCt^ method. At least three biological samples were analyzed to assess gene expression. A 95% Confidence Interval Estimation was applied, and values outside this range were excluded from the analysis.

Primers were designed to meet lengths ranging from 18 to 24 base pairs, melting temperature (Tm) around 60°C, amplicon size between 75 and 200 bp, GC content between 40% and 60%, and a low likelihood of forming hairpins and primer dimers. When possible, primers overlapping two exons were designed. Primer pair efficiency was verified with a standard curve and an efficiency of 90%-110% was required for RT-qPCR analysis.

### Phenotyping

30 to 40-day-old plants were collected for analysis of fresh weight, rosette area, and leaf number. Fresh weight was determined on a precision balance immediately after the collection of each rosette. The rosette area was then estimated using the Easy Leaf Area 2.0 tool (Easlon & Bloom, 2014), which relies on the color proportions of each pixel to distinguish the leaves and the calibration areas of the background (Easlon & Bloom, 2014). Finally, the number of leaves was counted, starting with the youngest and ending with the oldest.

### Starch Extraction and Quantification

Starch content was assessed in the insoluble fraction remaining after the ethanolic extraction of soluble sugars, followed by enzymatic digestion with α-amylase and α-glucosidase as described in Viana et al., 2021.

### Microarray

RNA profile analysis by microarray approach was essentially as described in Viana et al., 2021. Robust Multi-array Average (RMA) normalization was carried out using Expression Console™ (Affymetrix), and statistical analysis of the data was performed with the _AFFYLM_GUI R package (R Core Team, 2018). The *p* value *p<* 0.05 was used as the threshold for selecting differentially expressed genes and assessing expression levels between genotypes.

### Protein Extraction and Western Blot

Protein extraction was carried out using RIPA buffer (MERCK, R0278) containing One Pierce™ Protease Inhibitor Mini Tablet (ThermoScientific, 88665) per 10 mL. 500 µL of buffer was added to 15 mg of Arabidopsis leaf samples, which had been previously ground in liquid nitrogen. Samples were kept on ice for 5 minutes, homogenized every minute, and centrifuged at 13,000 rpm for 30 minutes at 4°C. The supernatant was transferred to Vivaspin® 500 3kDa concentrators (Sartorius), centrifuged at 13,000 rpm for 60 minutes at 4°C, and stored at -80°C. Protein quantification was performed using the Pierce™ BCA Protein Assay Kit following the instructions of the manufacturer. A linear regression correlating absorbance at 562 nm to bovine serum albumin concentrations was established to calculate extracted protein concentrations. Proteins were fractionated in denaturing SDS-PAGE with a 8% resolving gel and a 4% stacking. For Western blot analysis, transfer and antibody incubation steps were done as described by Mahmood and Yang, 2012. An Amersham Hybond P 0.45 PVDF membrane was activated by immersion in methanol for 30 seconds on each side. The primary antibodies used were monoclonal Anti-HA (H3663, Sigma-Aldrich) diluted 6:10,000 and Anti-Plant Actin (A0480, Merck) diluted 0.25:10,000. The secondary antibody was Anti-Mouse IgG (Goat Anti-Mouse IgG (H+L)-HRP Conjugate, 1721011, ThermoScientific), diluted 1:10,000. Detection was performed using the Amersham™ ECL™ Prime reagents. Chemiluminescence, resulting from HRP-mediated oxidation of luminol, was digitally captured. Densitometry analysis was conducted with ImageJ’s Gel Analysis tool. Data normalization was based on actin.

### Data Analysis and bioinformatics

For normally distributed data (Shapiro-Wilk and/or Levene tests, *p*>0.05), analyses were performed using Student’s t-test for pairwise comparisons and one-way ANOVA followed by Tukey’s post hoc test for multiple comparisons (*p*<0.05). For non-normally distributed data (Shapiro-Wilk and/or Levene tests, *p*<0.05), pairwise comparisons were conducted using the Wilcoxon test, and multiple comparisons were assessed using the Kruskal–Wallis test followed by Dunn’s post hoc test (*p* < 0.05).

Overlaps between gene lists were carried out using the webtool Venn Diagram (http://bioinformatics.psb.ugent.be/webtools/Venn/).

Heat maps, graphs and Statistics were performed with the R programming language (R 4.3.2, 2023, R Foundation) and figures were created using the vector graphic design software Inkscape (1.2.2, 2023, Inkscape Project).

## RESULTS

### *bZIP63* regulates the expression of subset of genes induced under short-day photoperiod

*bZIP63* is a key player in energy homeostasis and is essential to adjust growth and development with energy availability in synchrony with the diel cycle (Baena-González et al., 2008; Mair et al., 2015; Pedrotti et al., 2018; Frank et al., 2018; Viana et al., 2021). To further evaluate the role of *bZIP63* in energy management we expanded our previous analysis of End of Night (EN) *bzip63-2* T-DNA mutant leaf transcriptome (Viana et al., 2021) by exploring the End of Day (ED) leaf RNA profiles of *bzip63-2* (Ws background; Table S1). In addition, the EN leaf transcriptome of the *RNAiWs_L9* line (Ws background), in which *bZIP63* expression is silenced to less than 10% of wild type *bZIP63* mRNA level by an RNAi approach (Frank et al. 2018), was also obtained (Table S1).

The Comparison of the *bzip63-2* ED Differentially Expressed Genes (DEG; *p<*0.05 and ≥|1.5| fold change) with *bzip63-2* EN DEG (Viana et al., 2021) defined a Non-*R*edundant set of 187 Downregulated (NR *bzip63-2* - D) and 265 Upregulated (NR *bzip63-2* - Up) genes in *bzip63-2* as compared to the wild type (Table S1; Table S2.1). In *RNAiWs_L9*, 461 Down (*RNAiWs_L9* - D) and 371 Upregulated (*RNAiWs_L9* - Up) genes relative to the wild type (*p<*0.05 and ≥|1.5| fold change) were identified (Table S1; Table S2.2). A comparative analysis of the DEG in *bzip63-2* and *RNAiWs_L9* with the genes responding to the transient over expression of KIN10 (KIN10ox), one of the catalytic sub-units of SnRK1 (Baena-González et al., 2007) revealed a most significant overlap between NR *bzip63-2* - D, *RNAiWs_L9* - D genes and those induced by KIN10ox (166 genes; *p<* 1.4e^-158^; Figure 1A and Figure S1 for a complete comparative analysis). This result agrees with the role of *bZIP63* in SnRK1 signaling (Maier et al., 2015; Pedrotti et al. 2018; Viana et al, 2021) and we focused our further analysis on this collection of *RNAiWs_L9* and NR *bzip63-2* downregulated and KIN10ox upregulated genes.

**Figure 1.**
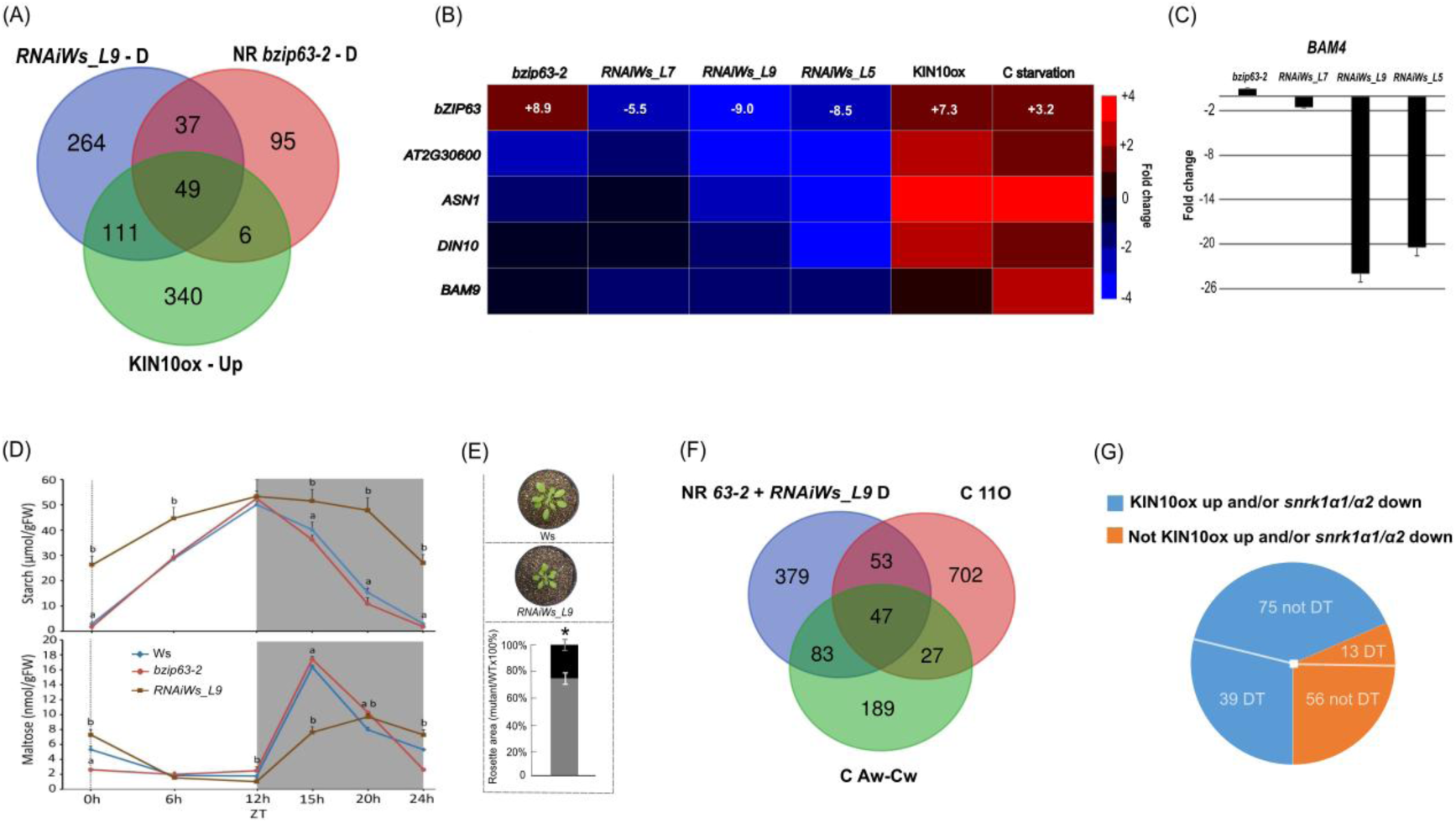
bZIP63 regulates a set of genes induced in short photoperiod. (A) Overlap of downregulated genes in *RNAiWs_L9* (*RNAiWs_L9* -D) and *bzip63-2* (downregulated at end of night and/or end of day, NR *bzip63-2* -D) with genes induced by KIN10ox (transient overexpression of a catalytic subunit of SnRK1; KIN10ox-Up; Baena-Gonzales et al., 2007). (B) Heat map Clustering of log2 fold changes relative to wild type of energy stress genes induced by KIN10ox and by carbon (C) starvation downregulated in *bzip63-2*, *RNAiWs_L7*, _*L9* and _*L5*. Gene expression was estimated in 30-day-old plants grown under short day condition (10h light /14 h dark) and 100 μmol m^−2^ s^−1^ of light intensity. All folds ≥ |1.5| are statistically significant compared to wild type (Student’s *t*-test; *p* < 0.05). (C) Fold change of *BAM4* expression in *bzip63-2* and *RNAiWs* lines *L7*, *L9*, and *L5* relative to Ws wild type. (D) Starch and maltose levels oscillation along the diel cycle (12h light / 12h dark) in *bzip63-2*, *RNAiWs_L9* line and WT rosette leaves of 30-day-old plants. The difference in starch and maltose levels between *bzip63-2* and *RNAiWs_L9* compared to WT was evaluated by one-way ANOVA with different letters indicating distinct groups (Tukey’s HSD test, *p* < 0.05). (E) Rosette area differences between wild type and *RNAiWs_L9* of 30-day old plants grown under 12h light / 12h dark and 100 μmol m^−2^ s^−1^ of light intensity. Statistical difference compared to wild type was determined by Student’s *t*-test (\**p* < 0.05). (F) Overlap between genes downregulated in NR *bzip63-2* and *RNAiWs_L9 (*NR *63-2 + RNAiWs_L9* D), short-day-induced genes that characterize the MDLM in short days (C Aw-Cw; Liu et al., 2021), and a set of 829 genes that are induced in the dark phase of short days (C 11O in Leung et al., 2023). (G) Categorization of the 183 downregulated genes shared between NR 63-2 + RNAiWs_L9 D, and the C Aw-Cw and C 11O clusters, based on whether they are upregulated in KIN10ox lines and/or downregulated in the *snrk1 α1/α2* mutant, and further distinguishing bZIP63 DT from non-targets.

A larger number of KIN10ox-induced genes were found to be downregulated in *RNAiWs_L9* (160 genes) than in *bzip63-2* (55 genes; Figure 1A). Moreover, among the 502 bZIP63-direct targets (DT) identified by Chromatin Immunoprecipitation (Muralidhara et al., 2021; Viana et al., 2021), 59 of them are specifically downregulated in *RNAiWs_L9*, whereas only five specifically overlap with NR *bzip63-2* – D and 28 are common to both *RNAiWs_L9* - D and NR *bzip63-2* - D (Table S2.2). For this latter set of 28 shared bZIP63-DT, the average down regulation is stronger in *RNAiWs_L9* than in *bzip63-2* (Table S2.2). The expression of four bZIP63-DT genes which are downregulated in *RNAiWs_L9* and NR *bzip63-2* and are induced by KIN10ox and carbon starvation (i.e., are therefore readouts for energy/carbon starvation; Figure S2), *AT5G30600*, *ASPARAGINE SYNTHETASE* (*ASN1*), *RAFFINOSE SYNTHASE* (*DIN10*) and *BETA AMYLASE 9* (*BAM9)* was evaluated in two other RNAi lines, *RNAWs_L5* and *_L7* (Figure 1B). The amount of *bZIP63* mRNA was reduced to 11% and 19 % of the wild-type level, in *RNAiWs_L5* and *RNAiWs_L7*, respectively (Figure 1B). This analysis showed a direct correlation between the extent of *bZIP63* silencing and the level of the downregulation of the four energy/carbon starvation readouts (Figure 1B). Moreover, the downregulation of these genes is stronger in the *RNAiWs_L9* and *RNAiWs_L5*, than in *bzip63-2* (Figure 1B). The reason for the differences between *bzip63-2* and *RNAiWs_L9* could partially be related to the strong downregulation of *BETA-AMYLASE 4* (*BAM4*) specifically in *RNAiWs_L9* (Figure 1C). *BAM4* encodes an inactive form of Beta-Amylase but acts as an activator of starch degradation (Fulton et al., 2008) and its downregulation in *RNAiWs_L9* resulted, as expected, in reduced starch decay and deregulation in maltose content relative to the wild type during the night (Figure 1D). This defect in starch utilization in *RNAiWs_L9* is likely to promote an energy/carbon starvation condition during the night period (Graff and al., 2010; Viana et al., 2021) which can explain the significant slower growth of *RNAiWs_L9* compared to the wild type (Figure 1E). This starvation condition may also contribute to the larger number of downregulated energy/C-starvation–related genes (i.e., those up-regulated by KIN10ox) in *RNAiWs_L9* compared to *bzip63-2* (Figure 1A).

One of the genes specifically downregulated in *RNAiWs_L9* is *PHLOEM PROTEIN 2-A13* (*PP2-A13, AT3G61060*; Supplementary table S2.1), a gene which is required for robust vegetative growth in short-day photoperiods (8 h light) and whose expression is controlled by the MDLM system (Liu et al., 2021; Wang et al., 2024). *PP2-A13* is part of a larger set of genes that show higher daily expression under short photoperiod versus long photoperiod (16 h light), mainly due to their induction during the dark phase (Liu et al., 2021). Recently, *PP2-A13* was also identified as part of an additional group of short-day–induced genes, named Cluster 11O (C 11O). This set includes genes that are highly expressed in darkness, generally leading to increased expression under short-day conditions (Leung et al., 2023).

We therefore compared the RNAiWs_L9-D and NR bzip63-2-D gene sets with a set of 1,101 genes that, like *PP2A-13*, are induced during the dark phase of short days (Figure 1F; Liu et al., 2021; Leung et al., 2023). This analysis revealed a significant overlap of 183 genes (*p*<1.568e^-117^) between the RNAiWs_L9-D/NR bzip63-2-D gene sets and the short-day–induced gene clusters (C Aw-Cw and/or C 11O; Figure 1F). Of these genes, 41% (75) display the same expression profile as *PP2A-13* (Cluster Aw in Liu et al., 2021; Cluster 11Oa in Leung et al., 2023). Nearly one third (52 genes) are classified as bZIP63-DT, and 62% (114 genes) are either induced in KIN10ox lines and/or downregulated in *snrk1 α1/α2 mutants* (Figure 1G; Tables S2.3 and S2.4; Baena-González et al., 2008; Pedrotti et al., 2018). Furthermore, 68% (124 genes) are under circadian clock control, and 88% (161 genes) exhibit diel oscillations in transcript abundance (Table S2.5; Figure S3). These data suggest that bZIP63 participates in the regulation of short-day–induced genes, partly through interacting with the circadian clock and in a SnRK1-dependent manner.

To further investigate the role of bZIP63 in the control of short-day-induced genes and to evaluate the reasons for the differences between *RNAiWs_L9* and *bzip63-2*, we produced and analyzed a *bzip63* CRISPR/Cas -induced frame shift mutant.

### CRISPR/Cas9-generated *bZIP63* frameshift mutation impairs growth but does not fully reproduce *RNAiWs_L9* characteristics

We obtained two independent CRISPR/Cas9-induced *bZIP63* frameshift mutant lines in the Col-0 background, designated *bzip63-5* and *bzip63-15* (Figure S4). Both lines exhibited a 60% reduction in *bZIP63* mRNA levels relative to wild-type plants and showed comparable deregulation of *AT2G30600* and *ASN1*, two direct targets of bZIP63 (Figure S5).

For further analysis, we focused on *bzip63-5*. We assessed, in short day photoperiod conditions (8h light / 16h dark), the expression of a defined set of short-day-induced genes (SDIG) involved in distinct biological processes: senescence (*SENESCENCE-ASSOCIATED GENE 1*, *SEN1*), amino acid metabolism (*ASN1*; *3-METHYLCROTONYL-COA CARBOXYLASE 1*, *MCCA*; *PROLINE DEHYDROGENASE 1*, *ProDH*), carbohydrate metabolism (*RAFFINOSE SYNTHASE 6*, *DIN10*; *MYO-INOSITOL OXYGENASE 2*, *MIOX2*), starch metabolism (*BETA-AMYLASE 4*, *BAM4)* and protein containing two BTB/POZ domains (*AT2G30600*). All of these genes are direct targets of bZIP63 (Muralidhara et al., 2021) and will be defined hereafter as bZIP63-related SDIG. Although *BETA-AMYLASE 9* (*BAM9*) is not classified as an SDIG, it is downregulated in bZIP63 mutants and has been identified as a direct target of bZIP63 (Viana et al., 2021) and was therefore included in the analyses as a marker of *bZIP63* activity. In addition, *PP2A-13*, which is not known to be a direct target of bZIP63 but is deregulated in *RNAiWs_L9* and serves as an MDLM marker gene under short-day conditions, was also analyzed (Figure 2). The expression of these genes was compared between *bzip63-5*, *bzip63-1* (a leaky mutant in the Col-0 background; Matiolli et al., 2012), *bzip63-2* and *RNAiWs_L9*. *bzip63-5* and *bzip63-2* exhibited similar patterns of gene expression deregulation, while *bzip63-1*, consistent with its leaky nature, showed a weaker response (Figure 2A). In contrast, *RNAiWs_L9* displayed the highest magnitude of gene expression deregulation, particularly for *BAM4*, consistent with the microarray data (Table S1). Thus, *bzip63-5* does not fully recapitulate the gene expression deregulation profile observed in *RNAiWs_L9* (Figure 2A).

**Figure 2.**
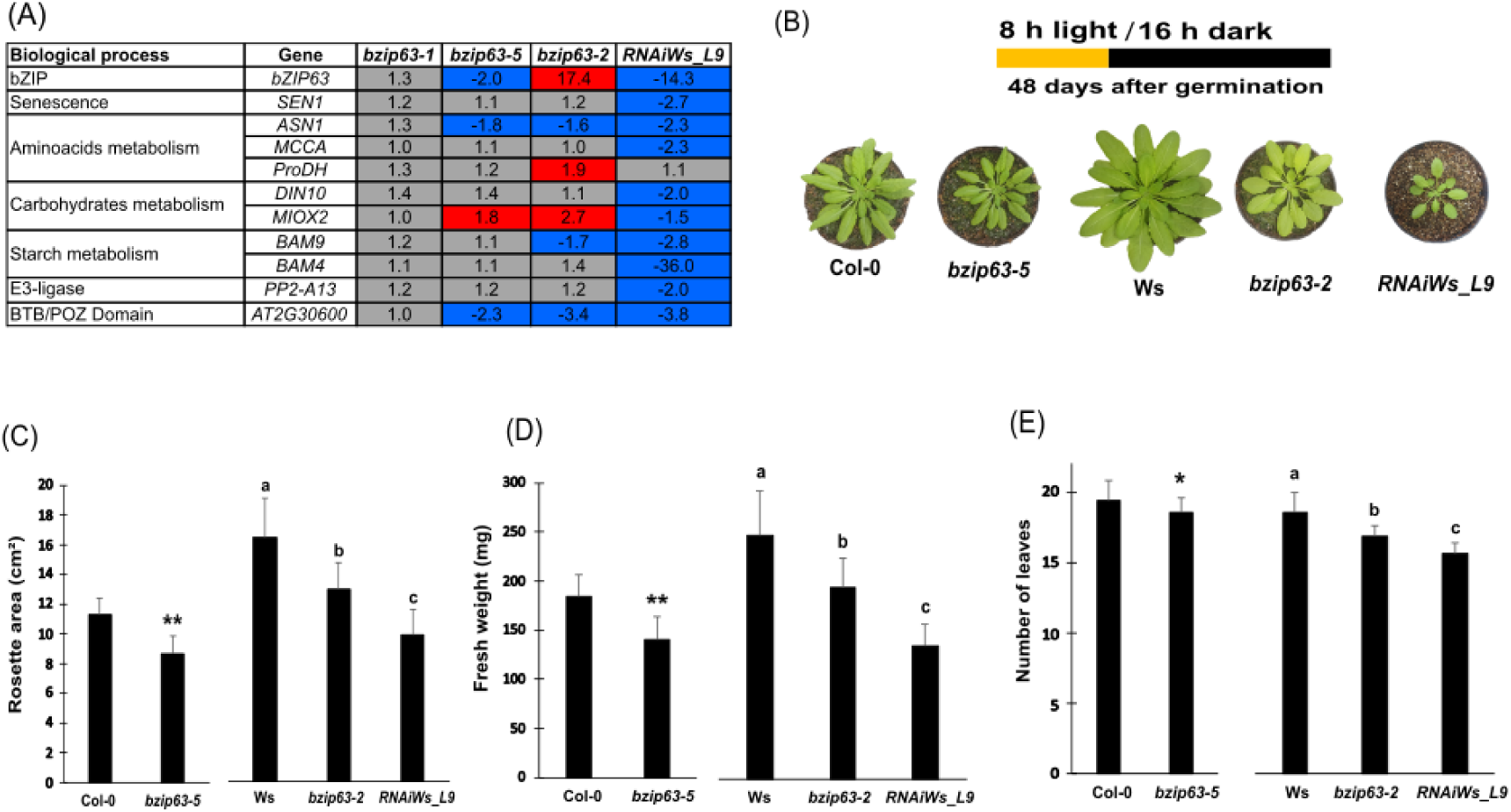
Arabidopsis *bZIP63* mutants exhibit reduced growth. (A) The CRISPR/Cas9-induced frameshift mutant line *bzip63-5* (Col-0 background) and the T-DNA insertion mutant *bzip63-2* (Ws background) similarly deregulate bZIP63-related SDIG, as well as *BAM9*, *BAM4* and *PP2-A13*, in leaves of 21-day-old plants sampled at the end of the night. Gray indicates fold change <|1.5|. Blue and red indicate downregulation and upregulation, respectively, for genes showing a fold change greater than |1.5| and a statistically significant difference compared to the wild type, as determined by Student’s *t*-test. (B) *bZIP63* mutants, Col-0 and Ws wild-types grown for 48 days after germination. (C) 40-day-old *bzip63-5*, *bzip63-2* and *RNAiWs_L9* plants exhibited growth impairment, characterized by smaller rosettes and reduced fresh weight (D) compared to wild-type plants. (E) The three mutant lines also showed a reduced number of leaves relative to wild type. Values are means and error bars indicate standard deviation. Statistical differences were assessed by Student’s *t*-test for Col-0 *vs bzip63-5* (*n* = 20, * *p*<0.05, ** *p*<0.01) and one-way ANOVA for Ws *vs bzip63-2 vs RNAi_L9*, with different letters indicating significantly distinct groups (Tukey’s HSD test, *p* < 0.05; n = 20). For leaf number (non-parametric), Wilcoxon was used for Col-0 *vs bzip63-5*, and Kruskal-Wallis with Dunn’s test for Ws, *bzip63-2*, and *RNAiWs_L9.* Plants were grown in short photoperiod (8 h light / 16 h dark), 80 μmol m^−2^ s^−1^ of light intensity.

We then compared the consequences of *bzip63-5*, *bzip63-2* and *RNAiWs_L9* mutations on vegetative growth, including rosette size, fresh weight and leaf number in short-day conditions. Both *bzip63-5* and *bzip63-2* mutants exhibited reduced rosette size compared to wild-type plants after 48 days of growth (Figure 2B). This reduction was already evident at 40 days, with decreases of 22% and 21% for *bzip63-5* and *bzip63-2*, respectively (Figure 2C). In contrast, the *RNAiWs_L9* line showed a more pronounced reduction of 40% relative to the Ws control (Figure 2B and C). These differences were correlated with fresh weight reductions of 24%, 21%, and 45% for *bzip63-5*, *bzip63-2*, and *RNAiWs_L9*, respectively (Figure 2D). The three mutant lines also developed fewer leaves than their respective wild types, further corroborating impaired growth (Figure 2E). Similar results were obtained in younger plants that were 30 days old (Figure S6).

To investigate the effect of photoperiod length on the growth of *bZIP63* mutants, we cultivated *bzip63-2* and *bzip63-5* under long-day conditions (16 h light / 8 h dark). To match the cumulative light exposure and equalize total light input from the short photoperiod experiments (8 h light / 16 h dark, 80 µmol m⁻² s⁻¹ for 30 days), plants were grown for 15 days under long-day condition with the same light intensity. Under these conditions, no significant differences in growth were observed between the *bZIP63* mutants and their respective wild types, indicating that long photoperiod circumvent the growth impairments observed under short photoperiod (Figure S7). Overall, these results reinforce that *bZIP63* is specifically required to sustain optimal growth under short-day conditions.

Given that C-group bZIP transcription factors, including *bZIP63*, form dimerization networks with S1-group members, and that the S1-group members *bZIP1* and *bZIP53* act as key metabolic regulators under energy deprivation (e.g. extended night) and control *ASN1*, *ProDH* and *MIOX2* expression (Wildenhain et al., 2025; Mair et al. 2015; Dröge-Laser and Weiste, 2018), we investigated the combined roles of bZIP1, bZIP53, and bZIP63 in regulating bZIP63-related SDIG and growth under short-day photoperiods. To this end, the triple mutant *bzip1-1/bzip53-1/bzip63-5* was generated by crossing *bzip63-5* with the null mutant *bzip1-1* (T-DNA insertion; Col-0 background; Dietrich et al., 2011; Figure S8A) and the leaky mutant *bzip53-1* (T-DNA insertion; Col-0 background; Dietrich et al., 2011; Figure S8B). *bzip1-1/bzip53-1/bzip63-5* was then compared with the *RNAiWs_L9* line (Figure S8C and D).

In *RNAiWs_L9*, transcript levels of bZIP63-related SDIG were downregulated, except for *ProDH* (Figure S8C). In contrast, transcript levels in the triple mutant were either unchanged or less affected, similar to the pattern observed in the single *bzip63-5* mutant (Figure S8C; Figure 2A). Notably, *BAM4* expression remained unaffected in the triple mutant (Figure S8C). These results suggest that the combined mutations in *bZIP1*, *bZIP53*, and *bZIP63* do not fully replicate the effects of *bZIP63* silencing achieved through RNAi-mediated repression in *RNAiWs_L9*. Growth analysis further supports this distinction as the triple mutant showed smaller reductions in rosette size (23%) and fresh weight (30%), whereas *RNAiWs_L9* exhibited more pronounced decreases of 59% in rosette size and 67% in fresh weight compared to wild-type plants (Figure S8D). Moreover, the growth of the triple mutant was only slightly reduced (not statistically significant) when compared to the single *bzip63-5* mutant (Figure S9B, C and D). Together, these results support the notion that *bZIP1*, *bZIP53*, and *bZIP63* act in a common pathway in which *bZIP63* has a dominant role.

To further investigate the contribution of bZIP63 transcript and protein abundance to the regulation of bZIP63-related SDIGs, we generated chimeric constructs encoding the bZIP63 cDNA fused to an HA (hemagglutinin) tag and Green Fluorescent Protein (GFP). These constructs were expressed either under the control of the constitutive CaMV 35S promoter (35S:HA:bZIP63:GFP) or under the native bZIP63 promoter (pbZIP63:HA:bZIP63:GFP), and *bzip63-2* transgenic lines were subsequently produced.

### Role of bZIP63 fusion protein abundance and diel oscillation in bZIP63-related SDIG expression profile

The oscillation profile of the 161 genes that are downregulated in *RNAiWs_L9* and/or *bzip63-2*, induced under SD conditions (e.g., *PP2-A13*) and exhibit diel oscillations in transcript abundance, are predominantly dark-phased (Table S2.5 and Figure S3), similar to the oscillation pattern of bZIP63 itself. However, how these expression profiles correlate with bZIP63 protein abundance remains poorly understood. To address this issue, we analyzed *bzip63-2* lines expressing *bZIP63* to different levels.

Chimeric genes encoding bZIP63 cDNA fused to an HA (hemagglutinin) tag and the Green Fluorescent Protein (GFP) under the control of the CaMV 35S promoter (*35S:HA:bZIP63:GFP*) or under the control of bZIP63 own promoter (*pbZIP63:HA:bZIP63:GFP*) were built and transferred into *bzip63-2* (Figure S10A). Two and three independent *bzip63-2* transgenic lines carrying *35S:HA:bZIP63:GFP* (*bZIP63-ox3*, *bZIP63-ox4*) or *pbZIP63:HA:bZIP63:GFP* (*bZIP63-co1, bZIP63-co2, bZIP63-co3*) constructs, respectively, and producing detectable levels of the ∼69 kDa bZIP63-fusion protein were selected (Figure S10B, C, D and E). The *bZIP63-ox3* and *bZIP63-ox4* lines displayed comparable levels of *HA:bZIP63:GFP* transcripts and fusion protein, as well as similar expression levels of the bZIP63-DT genes *AT2G30600* and *BAM9* (Figure 3A and Figure S10B and C). Thus, line *bZIP63-ox3* was chosen for further analysis. Lines *bZIP63-co1*, *bZIP63-co2*, and *bZIP63-co3* also exhibited comparable levels of *bZIP63* fusion transcripts and fusion protein, and the expression of *AT2G30600* and *BAM9* was essentially restored to wild-type levels at the EN in all three lines (Figure 3A; Figure S10D and E). We therefore selected lines *bZIP63-co1* and *bZIP63-co2* for subsequent analysis. In these lines, *HA:bZIP63:GFP* driven by the native *bZIP63* promoter reproduced the endogenous *bZIP63* mRNA diurnal oscillation, peaking at the EN and strongly declining toward the ED (450-fold in *bZIP63-co1* and 254-fold in *bZIP63-co2*, compared with a 50-fold decline in Ws; Figure 3B). In contrast, in line *bZIP63-ox3*, expression driven by the 35S promoter showed a smaller, 1.7-fold, reduction of fusion transcript levels at the ED versus EN (Figure 3B). Similar results were obtained for lines *bZIP63-co1* (∼470-fold difference) and *bZIP63-ox3* (7-fold) in an independent experiment (Figure S10F).

**Figure 3.**
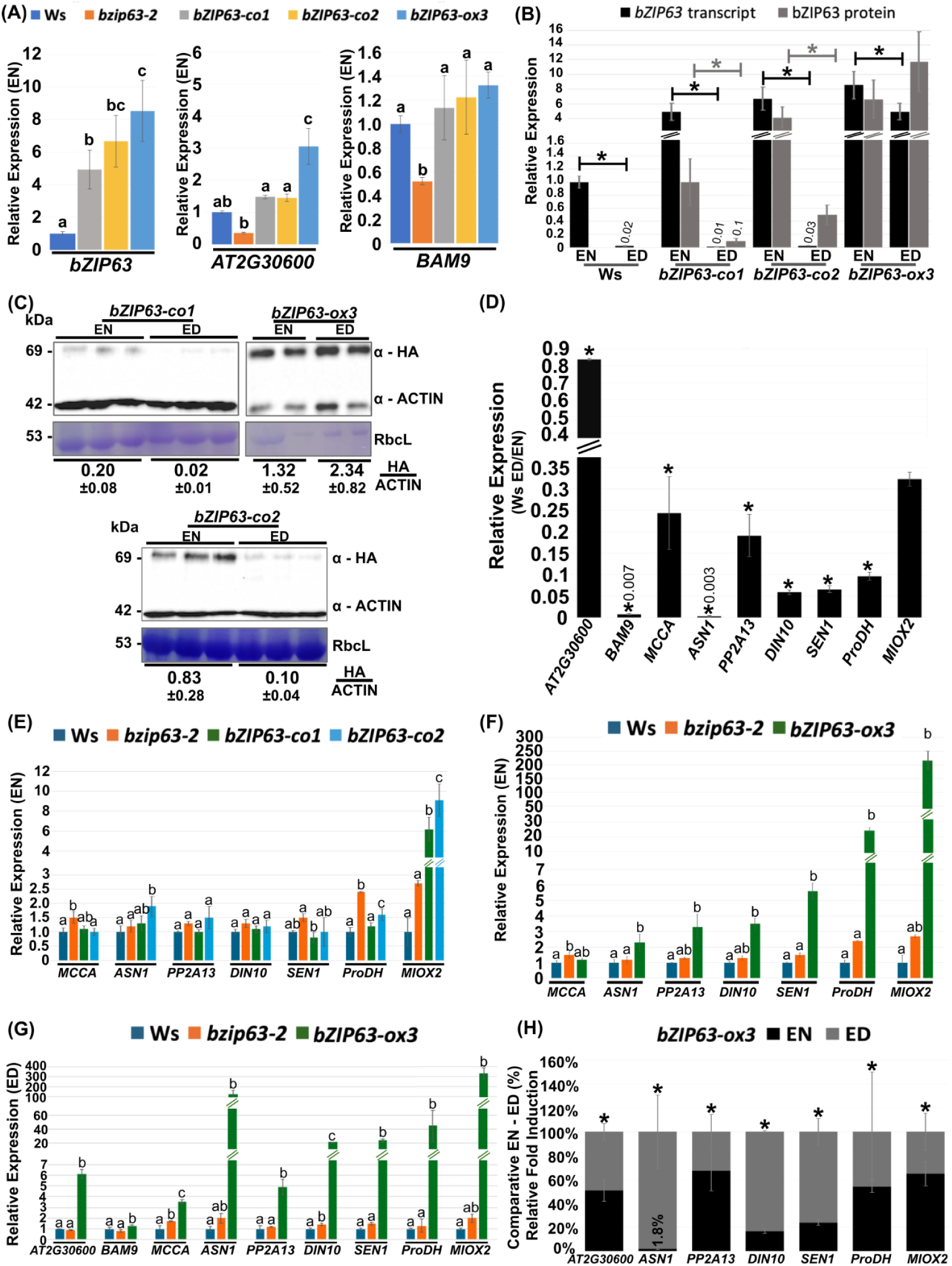
Diel oscillation in bZIP63 protein abundance and its influence on target gene expression. (A) Relative expression of *bZIP63* and its target genes *AT2G30600* and *BAM9* at EN. Expression values are relative to Ws. (B) Relative *bZIP63* mRNA levels, and abundance of bZIP63 protein in Ws, *bZIP63-co1*, *bZIP63-co2* and *bZIP63-ox3*. *bZIP63* transcript values are relative to Ws at the EN, and protein values represent relative abundance to *bZIP63-co1* at the EN. (C) Western blot analysis of HA:bZIP63:GFP accumulation in *bZIP63-co1*, *bZIP63-co2* and *bZIP63-ox3* at the ED and EN. The fusion protein has ∼69 kDa. Total protein loaded was 60 µg for *bZIP63-co1* and *bZIP63-co2*, and 5 µg for *bZIP63-ox3*. Actin (∼42 kDa) served as loading control and was used for normalization. Coomassie blue–stained gel shows Rubisco’s large subunit (RbcL, of ∼53 kDa). The average ratios and standard deviation of HA/ACTIN signals are indicated below the blots. (D) Relative expression of *bZIP63*-related SDIGs, *PP2-A13* and *BAM9* at ED in Ws. Values represent expressions relative to the expression of each gene in Ws at EN. (E) Relative expression of *bZIP63*-related SDIGs, *PP2-A13* and *BAM9* at EN in Ws, *bzip63-2*, *bZIP63-co1* and *bZIP63-co2*. Values represent expressions relative to the expression of each gene in Ws. (F) Expression of *bZIP63*-related SDIGs, *PP2-A13* and *BAM9* at EN in Ws, *bzip63-2* and *bZIP63-ox3*. Values represent expressions relative to the expression of each gene in Ws. (G) Relative expression of *bZIP63*-related SDIGs, *PP2-A13* and *BAM9* at ED in Ws, *bzip63-2* and *bZIP63-ox3*. Values represent expressions relative to the expression of each gene in Ws. (H) Fold induction of *bZIP63*-related SDIGs and *PP2-A13* in *bZIP63-ox3* relative to Ws at both EN and ED. The largest fold was set to 100% for each gene. All plants were grown for 21 days in short photoperiods and 80 μmol m^−2^ s^−1^ of light intensity. Values are means of two to three and three to five replicates per genotype for protein and transcript data, respectively, and error bars indicate standard deviation. Statistical differences in A, E, F and G were assessed by one-way ANOVA for parametric gene expression data, with different letters indicating significantly distinct groups (Tukey’s HSD test, p<0.05). For non-parametric gene expression data, the Kruskal-Wallis test was used, with different letters indicating significantly distinct groups (Dunn’s test, p<0.05). Statistical differences in B, D and H were assessed by Student’s t-test (* p<0.05).

In lines *bZIP63-co1* and *bZIP63-co2*, HA:bZIP63:GFP fusion protein levels accumulated at the EN and decreased approximately 10- and 8-fold, respectively, by the ED (Figure 3B and C). A similar trend was observed for line *bZIP63-co1* in an independent experiment (Figure S10F and G). In contrast, in *bZIP63-ox3*, fusion protein levels remained constitutively high at both EN and ED (Figure 3B and C; Figure S10F and G). The absolute levels of fusion protein in *bZIP63-ox3* at the EN reached approximately 13 and 8.5 folds the absolute levels in *bZIP63-co1* at EN in two independent experiments (Figure S10H and I). These data establish a positive correlation between fusion transcript abundance and fusion protein accumulation, and further suggest that, in the wild type, bZIP63 levels oscillate, peaking at the EN and reaching a trough at the ED.

These observations raise the possibility that the oscillatory pattern of bZIP63 protein abundance influences the oscillatory behavior of its target genes (Table S2.2), in agreement with previous proposals (Viana et al., 2021). To address this possibility, we examined the expression of the bZIP63-related SDIGs (*SEN1*, *ProDH*, *ASN1*, *AT2G30600*, *MIOX2*, *DIN10* and *MCCA*), *PP2-A13* and *BAM9* in lines *bZIP63-co1*, *bZIP63-co2*, and *bZIP63-ox3*, which differ in the abundance and oscillatory profile of the bZIP63 fusion protein (Figure 3B and C). These genes are dark phased, except for *AT2G30600* (Figure 3D).

In *bZIP63-co1* and *bZIP63-co2*, limited but significant changes were observed predominantly at the EN compared with the wild type and *bzip63-2* (Figure 3E; Figure S11 for ED data). The expressions of *AT2G30600*, *BAM9*, *MCCA*, and *ProDH* were mostly restored to wild-type levels (Figure 3A and E). *ProDH* and *MIOX2* transcript levels were higher in *bzip63-2* than in the wild type (Figure 3E; Figure 2A), consistent with negative regulation by bZIP63.

However, despite restoration of *bZIP63* expression, *MIOX2* was induced in *bZIP63-co1* and *bZIP63-co2* by approximately 6- and 9-fold, respectively (Figure 3E). In *bZIP63-ox3*, *MIOX2* expression was even more strongly induced, exceeding 200-fold at both the EN and ED relative to Ws (Figure 3F and G; Figure S12A and B). Similarly, *ProDH* expression in *bZIP63-ox3* was strongly increased by more than 20-fold at EN and ED compared to Ws (Figure 3F and G; Figure S12A and B). Together, these results indicate that both *MIOX2* and *ProDH* are subject to non-linear, bZIP63 dosage-dependent regulatory effects. Moreover, in *bZIP63-ox3*, *AT2G30600, ASN1, PP2-A13, DIN10,* and *SEN1* were significantly upregulated at both the EN and ED relative to Ws in two independent experiments (Figure 3A, F and G; Figure S12A and B). In contrast, *MCCA* and *BAM9* showed little or no deregulation (Figure 3A and F; Figure S12A and B).

*ASN1*, *DIN10* and *SEN1* tended to be more strongly induced at ED than at EN in *bZIP63-ox3* in two independent experiments (Figure 3H; Figure S13), suggesting a time-of-day–dependent competence to respond to bZIP63 overexpression. *PP2-A13*, *ProDH*, and *MIOX2* also tended to show stronger induction at ED (Figure 3H), however, this pattern was weaker and not reproducible across experiments (Figure 3H; Figure S13), indicating only a limited, if any, time-of-day dependence.

Finally, the EN-ED oscillatory pattern of *AT2G30600*, *ASN1*, *PP2-A13*, *DIN10*, *SEN1*, *ProDH*, *MIOX2* and *BAM9* transcript levels (Figure 3D) was maintained in *bZIP63-co1* and *-co2* and was partially maintained in *bZIP63-ox3* (Figure S14) despite the overall higher expression of these genes.

Together, these results indicate that the responsiveness of bZIP63-related SDIGs, *PP2-A13,* and *BAM9*, to changes in bZIP63 abundance differs markedly among them and is partially dependent on the time of day, consistent with gene-specific regulatory sensitivity, and may reflect a partial interaction with clock regulation of these genes.

## DISCUSSION

Seasonal photoperiod changes have profound effects on plant growth and development (Gendron and Staiger, 2023). Photoperiodic vegetative growth has recently been shown to be controlled by the MDLM system which evaluates the daily production of photosynthetic-derived sugar (photosynthetic period) to measure the day length. This system relies on circadian clock and photoperiodic control of sucrose and starch metabolism to promote photoperiod-specific gene expression pattern of *PP2-A13* and *MIPS1*, which are required for efficient SD and LD vegetative growth, respectively (Gendron and Staiger, 2023; Liu et al., 2021; Wang et al., 2024). *PP2-A13* exhibits a dark-phased expression pattern, with higher daily transcript levels in short days than in long days (Liu et al., 2021). This expression profile is shared by a cluster of genes defined here as short day–induced genes (SDIG; Liu et al., 2021; Leung et al., 2023). Whether the expression of all SDIGs is controlled by the MDLM system remains unclear; however, these genes are enriched in bZIP G-box (CACGTG) and AP2/ERF (AGCCGCC) binding sites (Leung et al., 2023), suggesting that they are integrated into related regulatory networks.

We show here that, among the SDIG set (1,101 genes; Figure 1F, Table S2.1; Liu et al., 2021 and Leung et al., 2023), a total of 183 genes, including *PP2-A13*, were down-regulated in *bzip63* mutant lines (*bzip63-2* and/or *RNAiWs_L9*; Figure 1F; Table S2.1). Of these 183, 114 (62%) are deregulated by KIN10 overexpression and/or in the a1a2 SnRK1 double mutant (Figure 1A and G; Table S2.3 and S2.4). Notably, approximately one third of this latter set (39 genes) were previously identified as bZIP63 DT (Figure 1G; Table S2.3 and S2.4;), consistent with regulation by the SnRK1–bZIP63 pathway and with the enrichment of carbon/energy-starvation–responsive genes within this group (Figure S2).

RNAi-induced silencing of *bZIP63* (∼10% of Ws levels) in *RNAiWs_L9* resulted in a larger number of downregulated genes among KIN10ox-induced genes, than in the T-DNA insertion mutant *bzip63-2* (Figure 1A), which is most likely a knockout (Maier et al., 2015; Viana et al., 2021). In addition, among the 39 SDIGs identified as bZIP63 DT and deregulated in KIN10ox and/or the a1a2 SnRK1 double mutant, 20 were specifically downregulated in *RNAiWs_L9*, whereas only 1 was specific to *bzip63-2* and 18 were shared by both genotypes (Table S2.2–S2.4). This apparent paradox may be explained by the strong downregulation of *BAM4* in *RNAiWs_L9*, a positive regulator of starch degradation (Figure 1C and D; Fulton et al., 2008), leading to slower starch degradation and an altered maltose accumulation profile (Figure 1D; David et al., 2021), which would reduce nocturnal carbon/energy availability (e.g., sucrose). However, the relative contributions of *bZIP63* silencing and *BAM4* downregulation cannot be distinguished from our data. The exclusive downregulation of the MDLM marker gene *PP2-A13* in *RNAiWs_L9* may also reflect the starch degradation defect, since *PP2-A13* promoter activity is altered in mutants defective in starch degradation (Liu et al., 2021). However, direct transcriptional regulation by bZIP63 can not be excluded, as DAP-seq analyses show that C/S1 bZIP heterodimers containing bZIP63 (bZIP1/bZIP63, bZIP2/bZIP63, and bZIP44/bZIP63) bind to the *PP2-A13* promoter (Liu et al., 2023).

The expression pattern in SD photoperiods of a subset of 7 of the 39 SDIGs that are bZIP63 DT(bZIP63-related SDIGs), plus *PP2-A13*, *BAM4* and *BAM9* (Figure 2A) was similar in the *bzip63-5* CRISPR/Cas frameshift mutant in Col-0 background and in *bzip63-2* in the Ws background, which shows no *BAM4* deregulation and fewer deregulated genes than in *RNAiWs_L9*. This observation further supports a role for *BAM4* in defining the *RNAiWs_L9*-specific set of deregulated genes. The reason for BAM4 downregulation in *RNAiWs_L9*, as well as in the independent *RNAiWs_L5* line, remains unclear. The more pronounced reduction in growth characteristics (i.e., rosette area, fresh weight, and leaf number) observed in *RNAiWs_L9* compared with *bzip63-2* and *bzip63-5* under short-day conditions, relative to their respective wild types (Figure 2B–E), may be related to the reduce nocturnal carbon/energy availability due to starch degradation defect in *RNAiWs_L9*. The growth defects of *bzip63-2* and *bzip63-5* were largely reverted under long-day conditions, consistent with a short-day–specific requirement for *bZIP63*, possibly to compensate for reduced nocturnal energy availability relative to long-day conditions (Graf et al., 2010; Sulpice et al., 2014; Viana et al., 2021).

The involvement of the heterodimerization network among bZIP63, bZIP1, and bZIP53 in energy-deprivation responses and in regulating *ASN1*, *ProDH*, and *MIOX2* expression (Wildenhain et al., 2025; Mair et al., 2015; Dröge-Laser and Weiste, 2018), prompted us to compare the expression of bZIP63-related SDIGs, *PP2-A13*, *BAM4*, and *BAM9*, as well as growth phenotypes, between the *bzip1-1/bzip53-1/bzip63-5* triple mutant and *RNAiWs_L9*. Gene expression patterns of the triple mutant were similar to those of the single mutant *bzip63-5* (Compare Figure 2A and S8), and the defect growth phenotypes were only slightly more pronounced than in *bzip63-5* (Figure S9). Together, these data indicate that *bZIP63* plays a significant role in optimizing growth under short-day conditions, whereas bZIP1 and bZIP53 contribute only marginally and may be more relevant under stress conditions (Wildenhain et al., 2025).

Among the 39 SDIGs identified as bZIP63 DT and deregulated in KIN10ox and/or the a1a2 SnRK1 double mutant (Figure 1G), 34 (87%) display diel oscillations, of which 28 (72%) peaked during the dark phase in equinoctial light conditions, similar to *bZIP63* itself. (Tables S2.3–S2.5; Figure S3) and for a subset of them in SD (Figure 3D). These observations prompted us to examine the regulation of bZIP63 protein abundance to determine whether and how it could contribute to the phasing of bZIP63 DT gene expression. In two independent *bzip63-2* transgenic lines expressing the HA:bZIP63:GFP fusion protein from the native *bZIP63* promoter (*bZIP63-co1* and *bZIP63-co2*) and grown under SD conditions, expression of *AT2G30600*, *BAM9*, *MCCA*, and *ProDH* was mostly restored to wild-type levels indicating partial complementation (Figure 3A and E). In both lines bZIP63-co1 and -2 *HA:bZIP63:GFP* transcript levels peaked at EN and declined sharply at ED, closely resembling the oscillation pattern of bZIP63 mRNA in the wild type (Figure 3B). Furthermore, fusion mRNA and protein levels were closely correlated (Figure 3B and C; Figure S10). These observations suggest that bZIP63 protein in the wild type oscillates in phase with its transcript, consistent with degradation during the day of protein synthesized during the night. This conclusion was further supported by the finding that expression of *HA:bZIP63:GFP* from the heterologous CaMV 35S promoter, which minimizes diel oscillation of the fusion transcript, resulted in constitutive accumulation of the fusion protein (Figure 3B and C; Figure S10F and G). Hence, we propose that the diel fluctuation of bZIP63 levels results from the combined effects of transcriptional regulation by the circadian clock and carbon/energy status (Viana et al., 2021), mRNA stability control (Matiolli et al., 2012; Viana et al., 2021), and protein turnover. The resulting timing and amplitude of bZIP63 accumulation may therefore contribute to shaping the expression profile of its DT SDIGs. Consistent with this hypothesis, *At2g30600*, *ASN1*, *SEN1*, *DIN10*, *MIOX2*, *ProDH*, and *PP2-A13* are strongly induced, albeit to different extents, at both EN and ED in the *bZIP63-Ox3* line, which constitutively accumulates bZIP63, while their diel oscillation is maintained (Figure 3F and G; Figure S12A and B; Figure S13; Figure S14). Furthermore, the induction of *ASN1*, *DIN10*, and *SEN1* in *bZIP63-ox3* is more pronounced at ED relative to EN than for the other genes, underscoring a potential time-of-day–dependent regulatory role of bZIP63. The variable responsiveness of bZIP63-related SDIGs to different bZIP63 levels and time of day may reflect differences in promoter architecture that determine their sensitivity to bZIP63 and their integration into distinct regulatory networks, including circadian clock regulation (Viana et al., 2021). This is illustrated by the dose-dependent antagonistic regulation of *ProDH* and *MIOX2* (Figure 3E, F and G; Figure S12A). Both genes are repressed by bZIP63, as indicated by their upregulation in the *bzip63-5* mutant, but are strongly induced in the *bZIP63-ox3* line and, in the case of *MIOX2*, also in *bZIP63-co-2* and *-3* (Figure 3E; Figure S14).

The influence of SnRK1 on the expression of bZIP63-related SDIGs, including *PP2-A13* and *BAM9* (Figure 1G; Tables S2.3 and S2.4), together with the reported increase in SnRK1 activity in vivo toward EN and its maintenance during the day (Avidan et al., 2023), suggests that SnRK1-mediated phosphorylation of bZIP63 also contributes to the regulation of SDIG expression.

## Acknowledgements

This work was supported by the São Paulo Research Foundation (FAPESP) (grant nos. 2020/00304-3, 21/05512-6 and 23/11182-4), 2018/25710-4 (FAPESP/UKRI-BBSRC cooperation agreements), 2019/25993-9 (FAPESP/Max Planck Institute cooperation agreements) and Conselho Nacional de Desenvolvimento Cientıfico e Tecnologico (CNPq)/Cofecub (Grant 0043/2020).

## Author contributions

RAC and PTC designed the experiments and collected and analyzed gene expressions, protein, and phenotype data. AJCV performed starch/maltose quantification and conducted microarray analyses. IS performed and analyzed part of the gene expression experiments. JGPV, AJCV and CCM generated the bZP63 constructs and transgenic lines. MV designed experiments, analyzed data and provided funding. RAC, PTC, and MV wrote the manuscript. RAC and PTC contributed equally to this work.

## SUPPLEMENTARY MATERIAL

**Figure S1.**
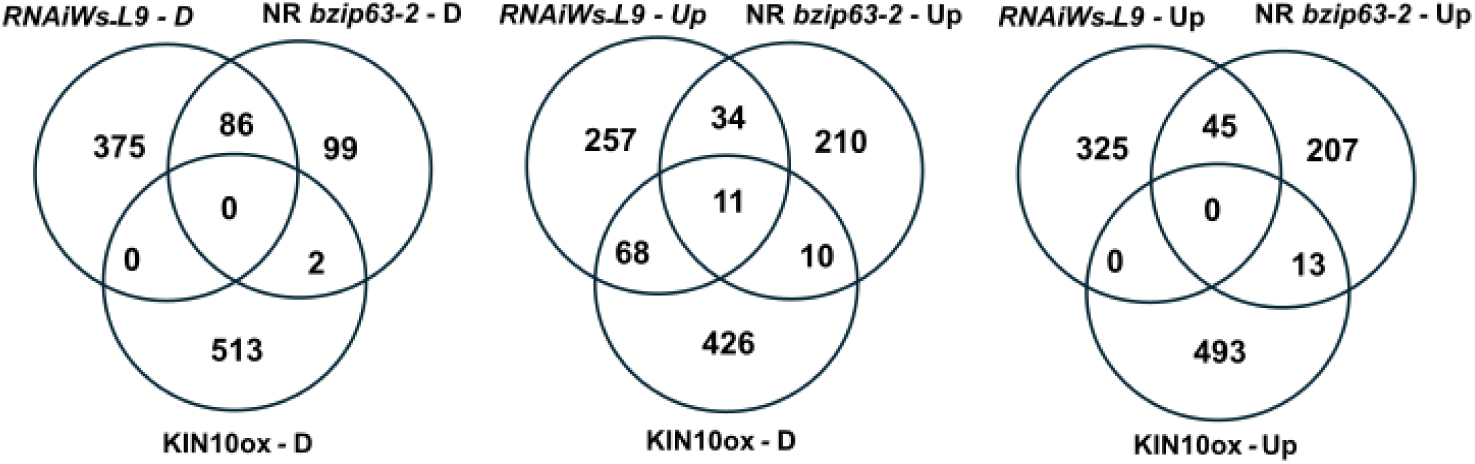
Venn Diagrams of the number of overlapping genes between Up or Down (D) regulated genes in *RNAiWs_L9* at the end of night (*RNAiWs_L9* - Up, *RNAiWs_L9* - D), in *bzip63-2* at the end of the night and/or end of day (NR *bzip63-2* - D, NR *bzip63-2* - Up) and those up or downregulated by overexpressing KIN10 (KIN10ox - D or Up). The overlap in *RNAiWs_L9* – Up / NR *bzip63-2* – Up / KIN10ox - D is significant (*p*< 1.195e^-54^).

**Figure S2.**
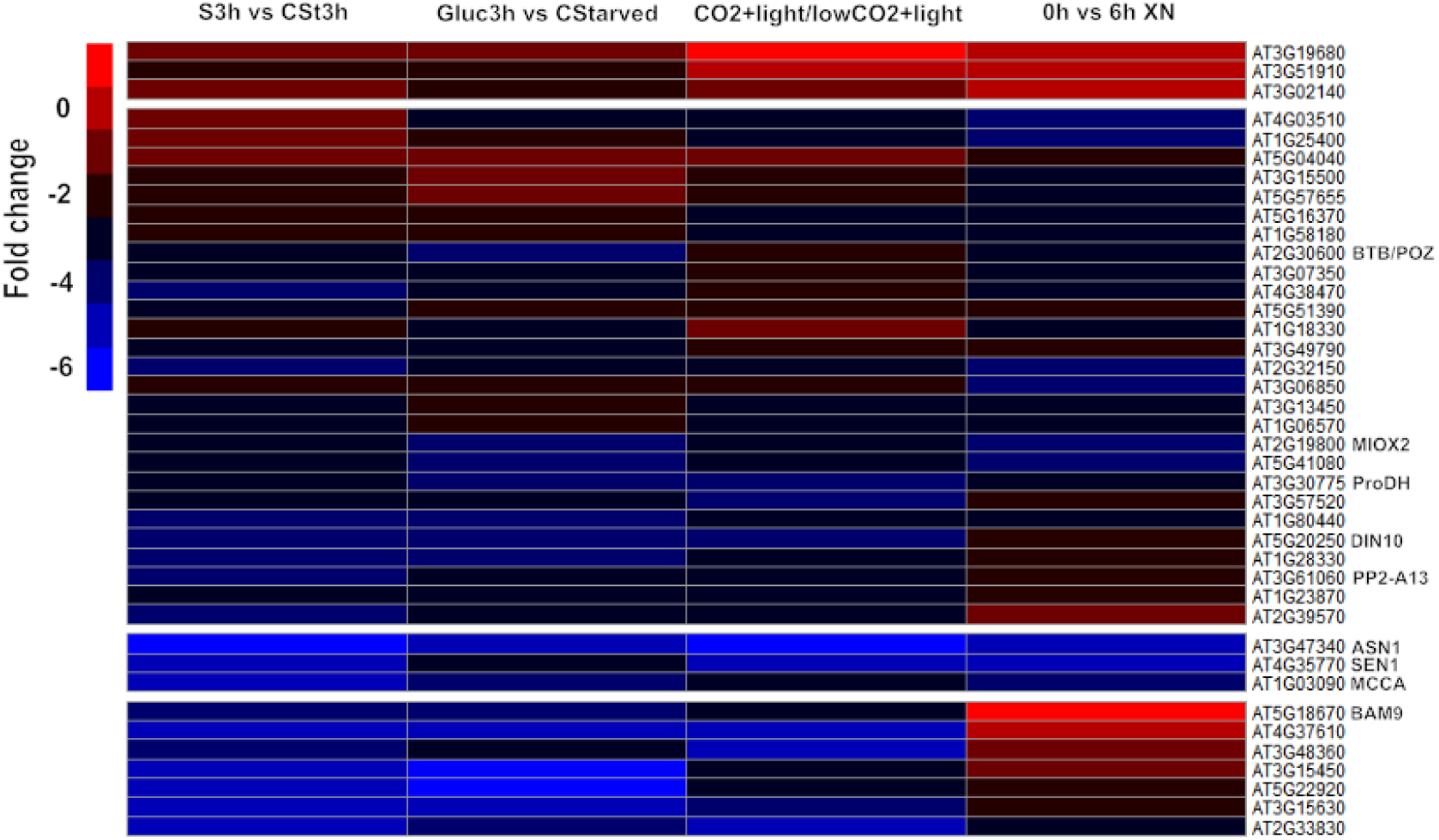
Mean transcript changes of short-day–induced bZIP63-target genes, as well as *BAM9* and *PP2-A13*, which are deregulated in KIN10ox or *snrk1α1/α2*, following sucrose, glucose, CO₂, and dark-extension treatments. S3h vs CSt3h, response 3 hours after addition of 15 mM sucrose to C-starved seedlings compared to C-starved seedlings (Bläsing *et al*., 2005); Gluc3h vs CStarved, response after 3 h adding of 100 mM glucose to C-starved seedlings compared to C-starved seedlings (Bläsing *et al*., 2005); CO2+light/lowCO2+light, the difference between rosettes illuminated at the end of the night for 4 hours at 350 or 50 ppm. [CO_2_] (Bläsing *et al*., 2005); 0h vs 6h XN, the difference between the end of the night and a 6-h extension of the night in wild-type plants (Thimm *et al*., 2004). All data are from Osuna et al., 2006.

**Figure S3.**
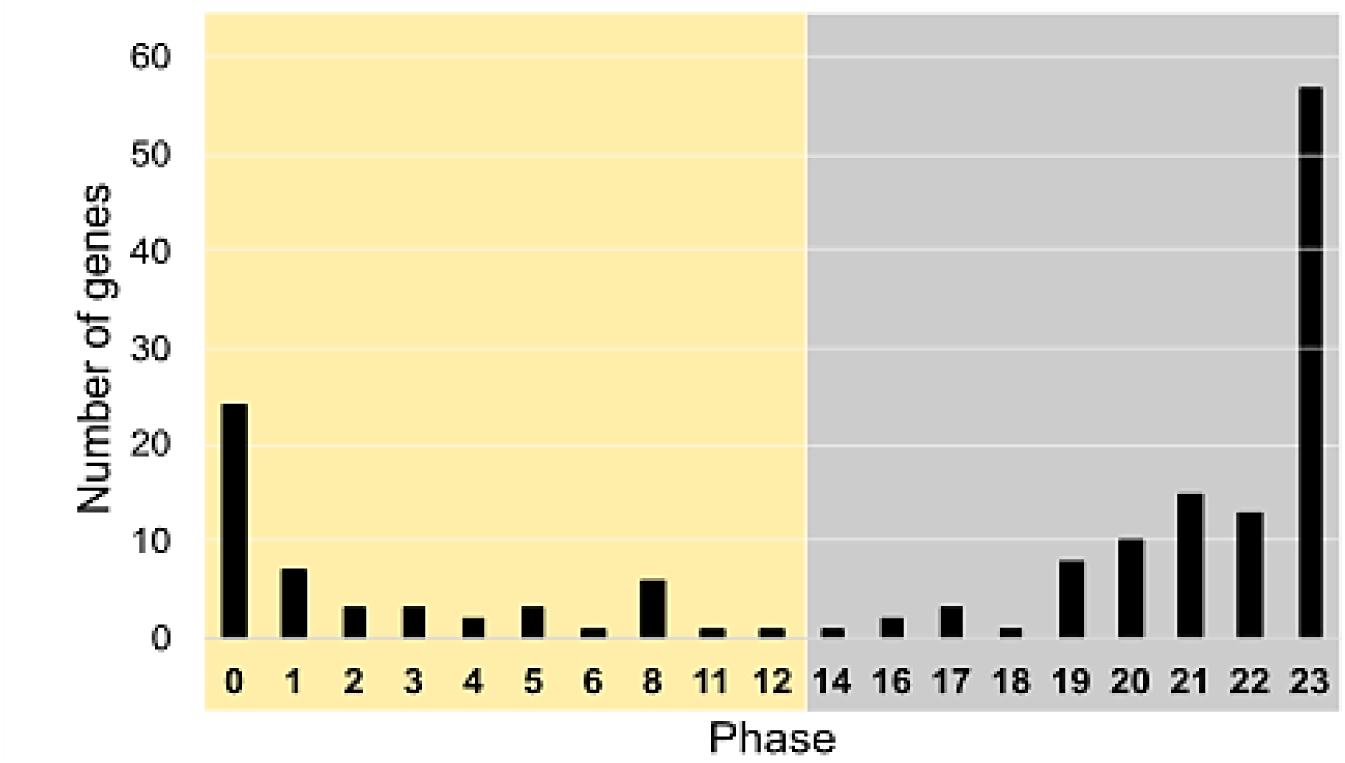
Diel oscillation of Cluster Aw-Cw and C11O genes downregulated in *bzip63-2* and/or *RNAiWs_L9*. Phase distribution of 161 genes from the NR63-2 + RNAiWs_L9_D gene set that overlaps with Cluster Aw-Cw (Liu et al., 2021) and C11O (Leung et al., 2023), exhibiting diel oscillations in mRNA abundance. Data were retrieved from the Diurnal web server (http://diurnal.mocklerlab.org/) and correspond to data from Bläsing et al., 2005.

**Figure S4.**
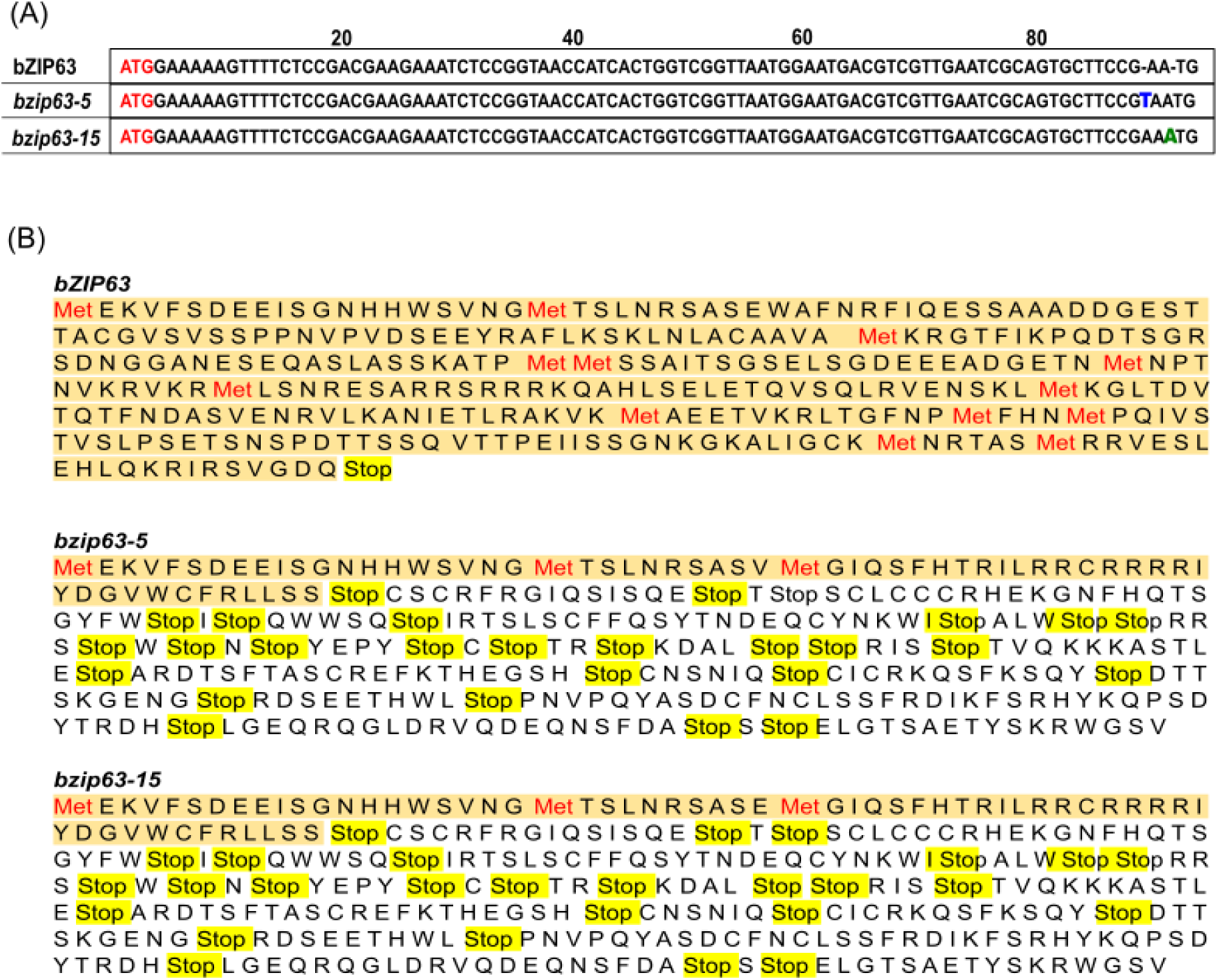
CRISPR/Cas9-induced frameshift mutations in *bZIP63* result in premature stop codons in *bzip63-5* and *bzip63-15* alleles. (A) Schematic representation of thymine (T) and adenine (A) nucleotide insertions into the coding sequence of the wild-type *bZIP63* allele, corresponding to the *bzip63-5* and *bzip63-15* genotypes, respectively. (B) The indels caused a frameshift mutation, leading to the formation of premature stop codons in both mutant alleles relative to the wild-type (Col-0) background.

**Figure S5.**
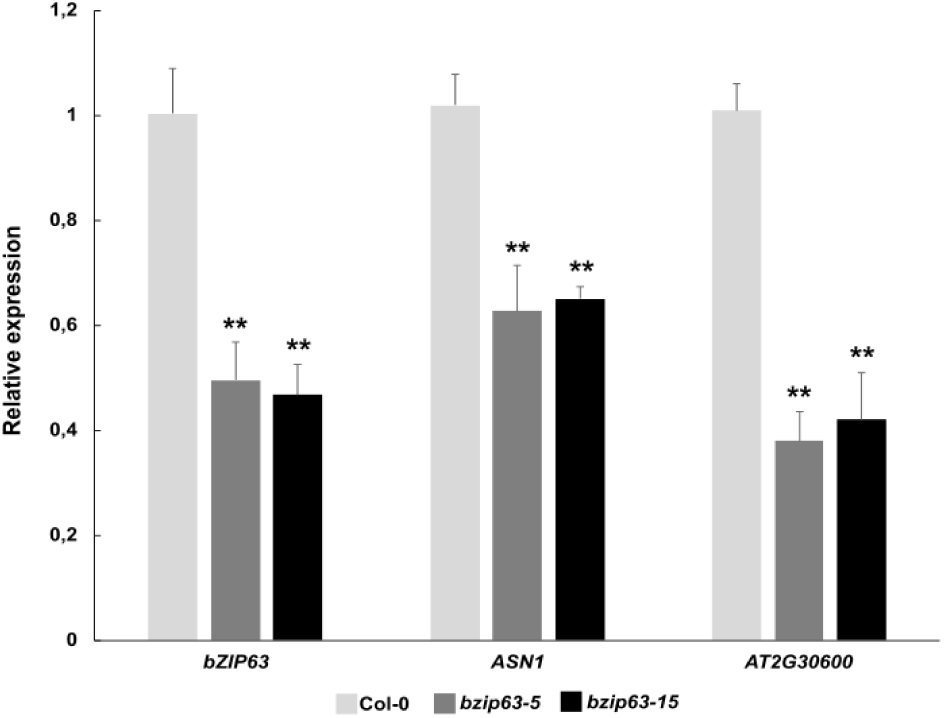
Characterization of *bzip63-5* and *bzip63-15* mutants. CRISPR/Cas-induced frameshift mutation lines, *bzip63-5* and *bzip63-15* (Col-0), retained 40% of wild type *bZIP63* mRNA level and exhibit identical levels of down regulation of *AT2G30600* and *ASN1*, two *bZIP63* activity readout genes. Values represent means, and error bars indicate standard deviation (Student’s *t*-test; \**p*<0.05; \*\**p*<0.01 compared to WT). Plants were grown 25 days after germination in short photoperiod, light intensity of 80 μmol m^−2^ s^−1^ and sampling was at the end of night.

**Figure S6.**
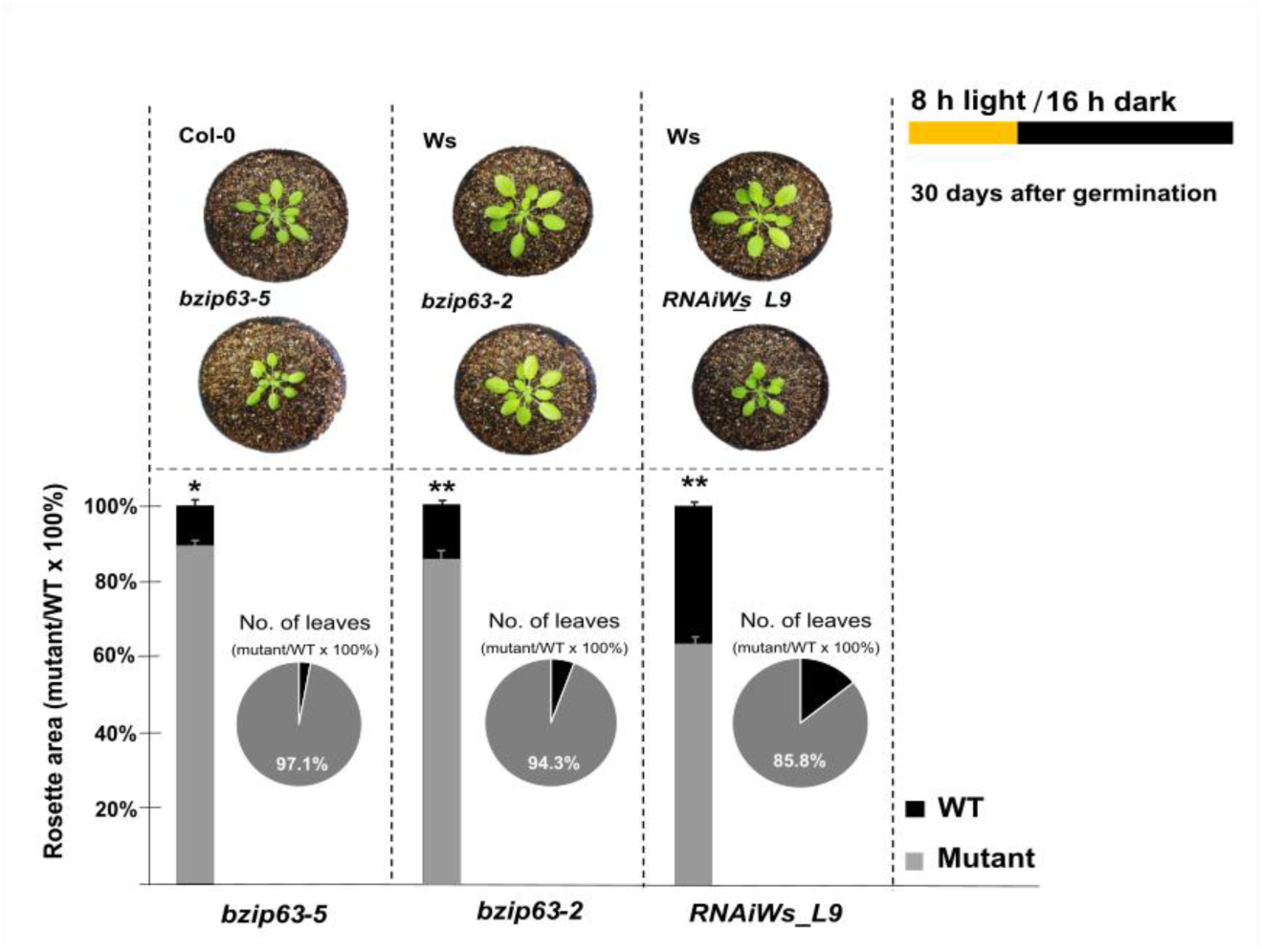
Reduced rosette size and leaf number in *bzip63* mutants. 30-day-old *bzip63-5*, *bzip63-2*, and *RNAiWs_L9* plants grown under short photoperiod conditions and exposed to a light intensity of 80 μmol m^−2^ s^−1^ exhibited impaired growth compared to wild-type (WT) plants (Student’s *t*-test; n = 20; \**p*<0.05, \*\**p*<0.01 vs. WT). Data are presented as means ± standard deviation. For reference, 100% leaf number corresponds to 14 leaves in Col-0 and Ws ecotypes, while 100% rosette area corresponds to 6 cm² in Col-0 and 9 cm² in Ws.

**Figure S7.**
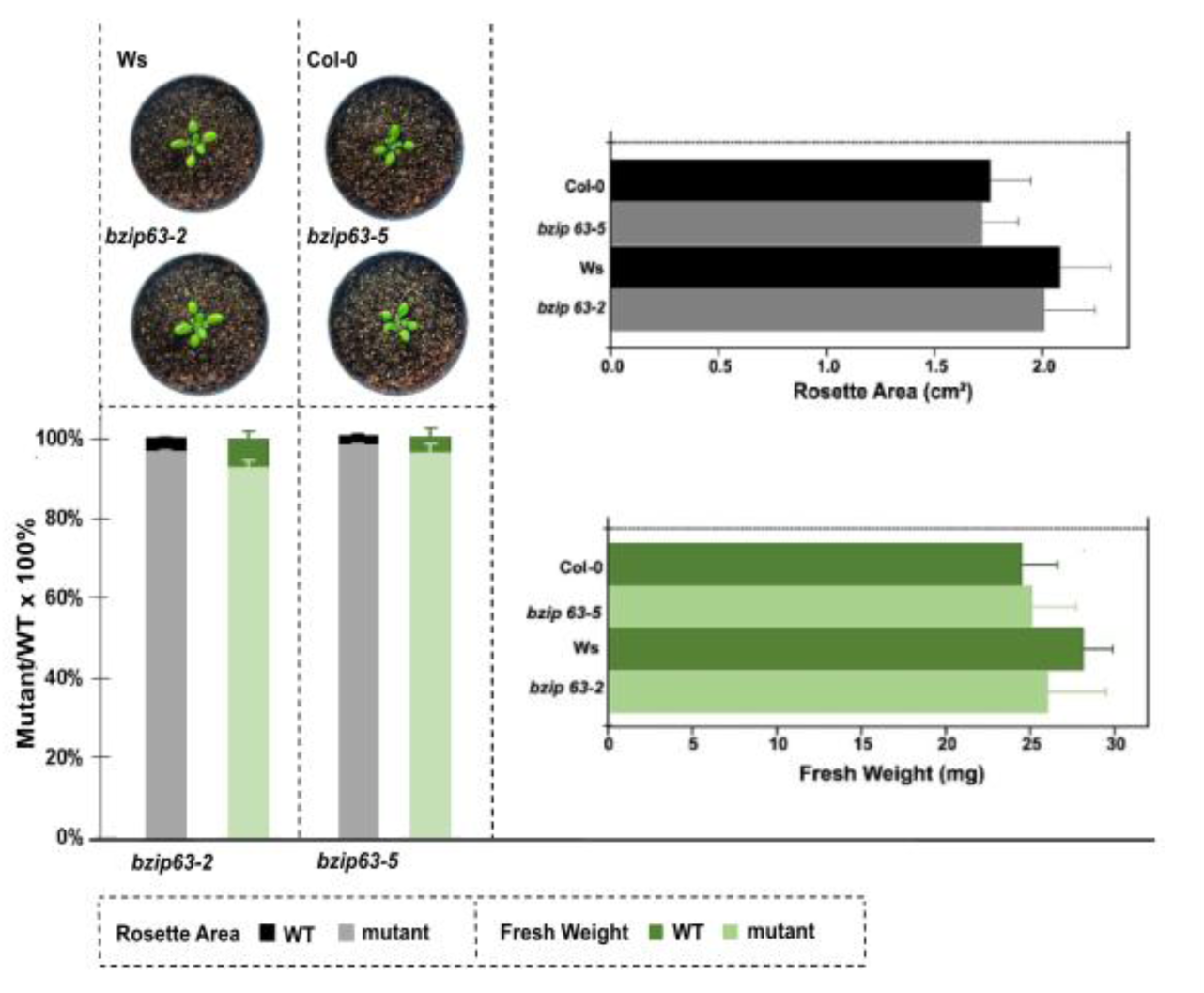
*bzip63-5* and *bzip63-2* exhibit wild type growth under long photoperiod. 15-day-old *bzip63-5* and *bzip63-2* mutants, along with Col-0 and Ws wild-type plants, were grown under long-day conditions (16 h light / 8 h dark) at a light intensity of 80 µmol m⁻² s⁻¹. Data are presented as means ± standard deviation. Statistical analysis was performed using Student’s *t*-test (n = 20; *p*<0.05, *p*<0.01 vs. WT).

**Figure S8.**
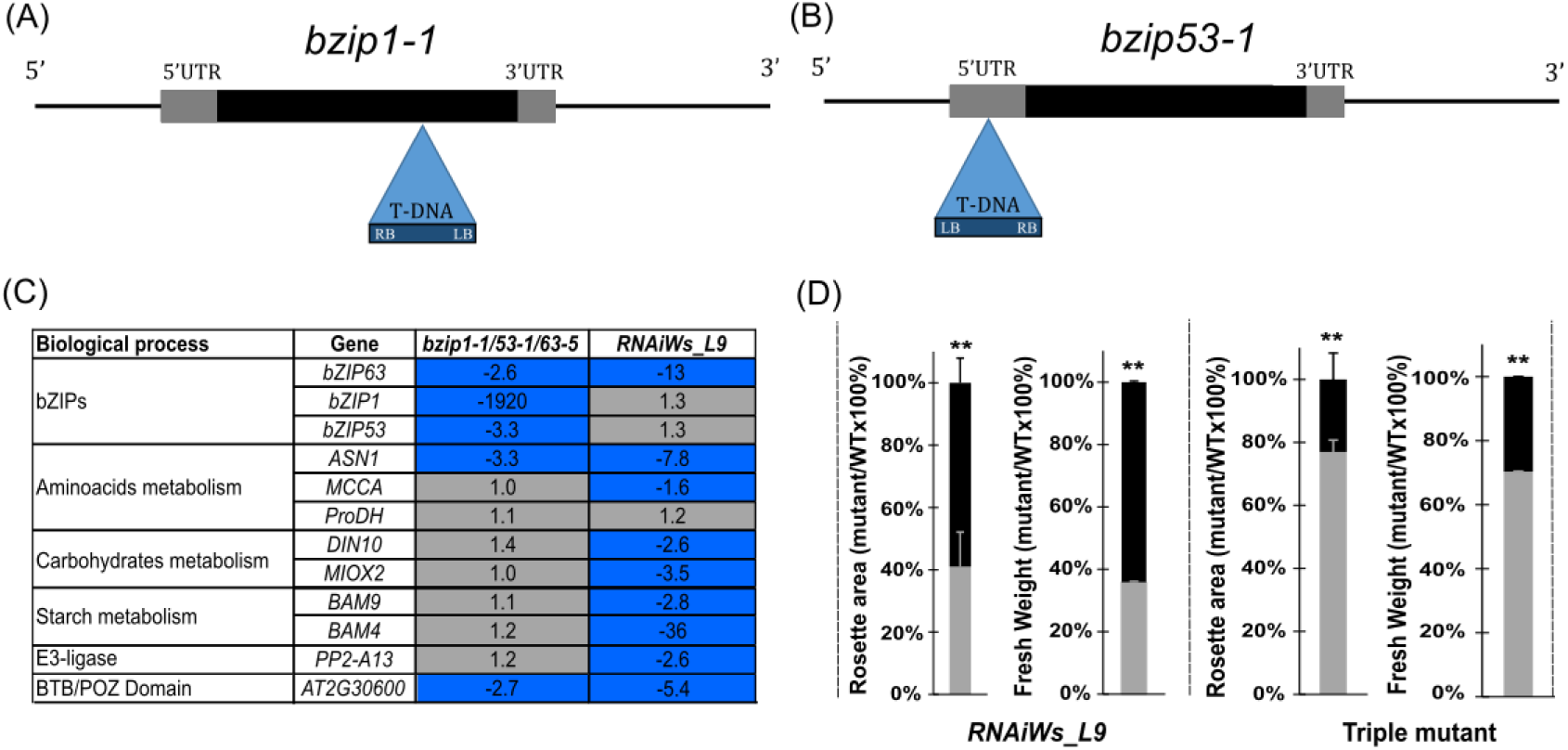
Characterization of the *bzip1-1/bzip53-1/bzip63-5* triple mutant and its impact on bZIP63-related SDIG expression and growth under short-day conditions. (A) The null mutant *bzip1-1* and the leaky mutant *bzip53-1* have a T-DNA insertion in *bZIP1* exon and *bZIP53* 5’- UTR, respectively (Dietrich et al., 2011). LB denotes the left border of the T-DNA, and RB denotes the right border. (C) Effects on transcript levels of bZIP63-related SDIG and *BAM9* in the triple mutant versus *RNAiWs_L9* in 30-day-old plants. Gray color indicates fold change <|1.5| and blue color indicate downregulation with fold changes ≥ |1.5| and statistical significance (Student’s *t*-test; *p*<0.05) compared to WT. (D) At 42 days, both the triple mutant and *RNAiWs_L9* showed growth impairment, characterized by smaller rosettes and reduced fresh weight compared to wild-type plants (Student’s *t*-test; n = 20; \*\**p*<0.01 vs. WT). Plants were grown in 100 μmol m^−2^ s^−1^ of light intensity. Values represent means, and error bars indicate standard deviation.

**Figure S9.**
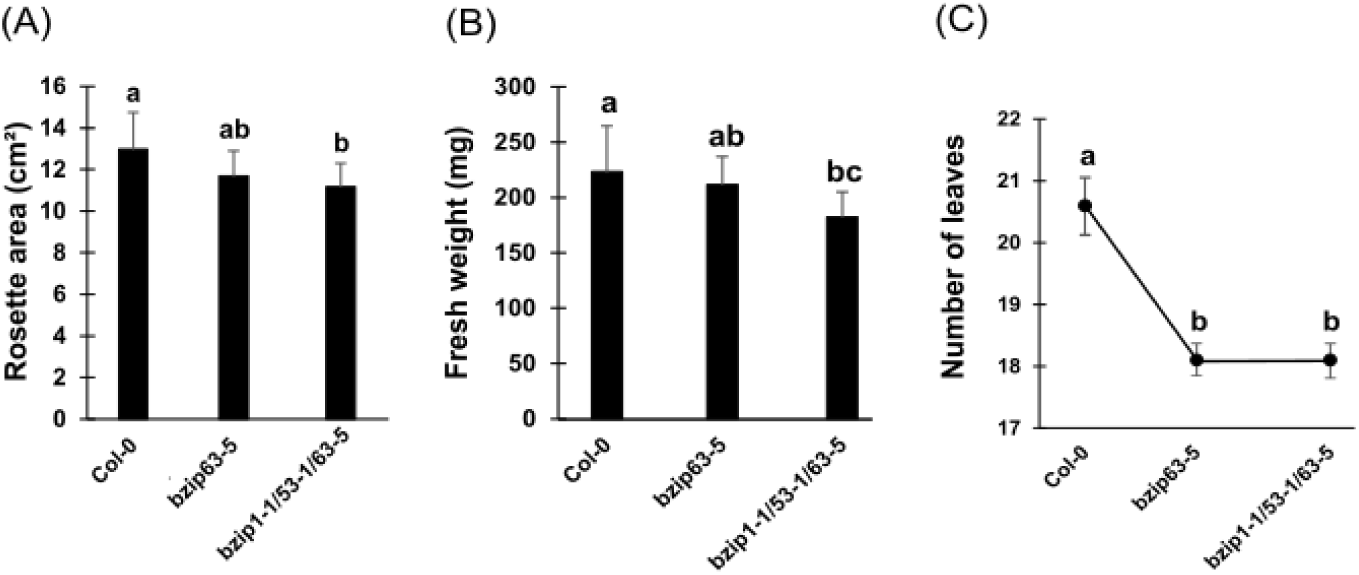
Triple *bZIP1/53/63* mutations do not exacerbate growth defects beyond *bZIP63* mutation alone. (A) Forty-day-old *bzip63-5* and triple mutant (*bzip1-1/bzip53-1/bzip63-5*) plants grown in 80 μmol m^−2^ s^−1^ of light intensity exhibited reduced rosette size and fresh weight (B), as well as fewer leaves (C), compared to wild-type plants. Data represents means, with error bars indicating standard deviation. Statistical differences were assessed by one-way ANOVA followed by Tukey’s HSD test (*p*<0.05; n = 20) for rosette area and fresh weight, with different letters indicating significantly distinct groups. For leaf number (non-parametric), Kruskal-Wallis with Dunn’s test was applied.

**Figure S10.**
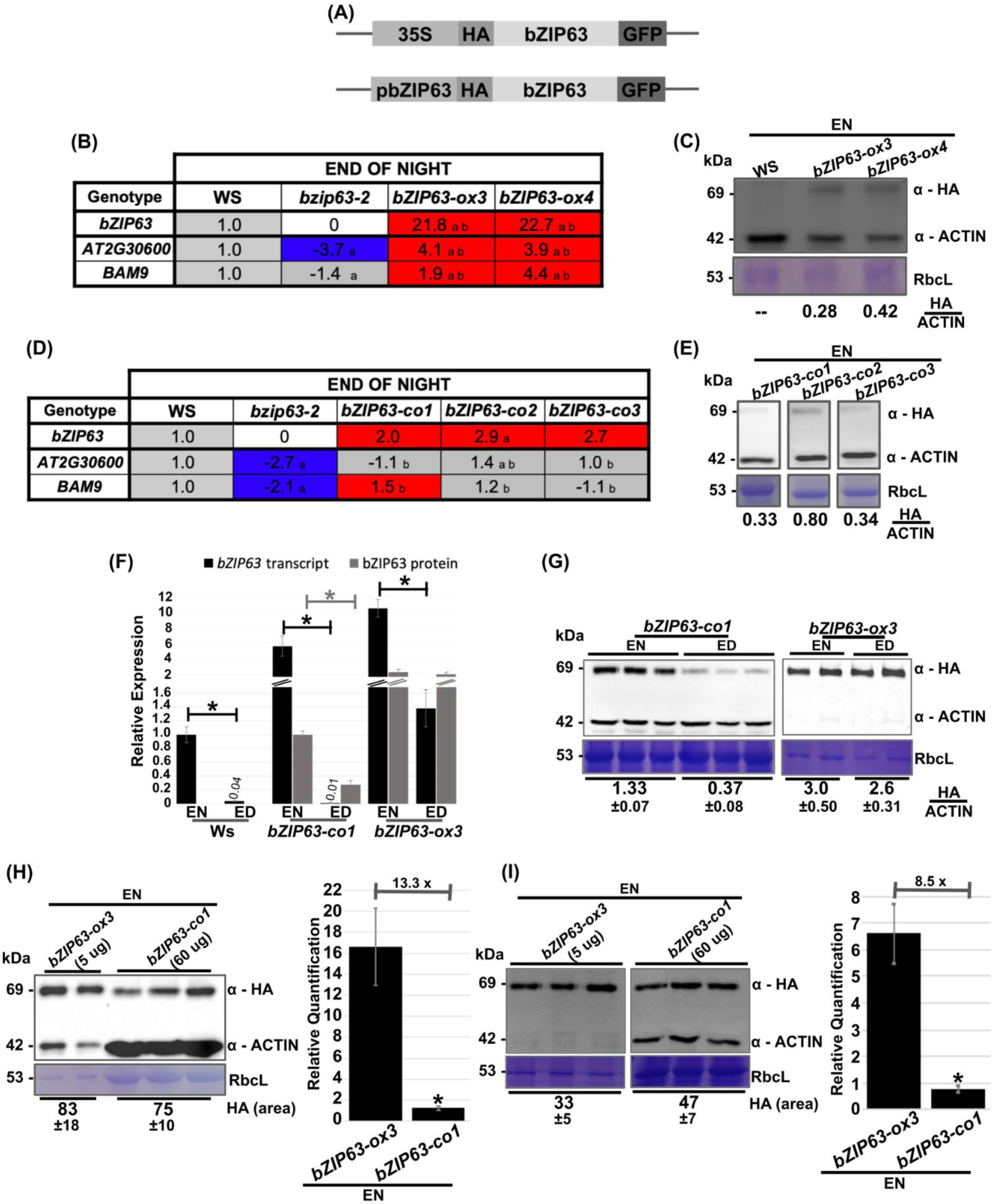
Characterization of *bzip63-2* lines expressing the HA:bZIP63:GFP construct (*bZIP63-ox3*, *bZIP63-ox4*, *bZIP63-co1*, *bZIP63-co2* and *bZIP63-co3*). (A) Schematic representation of constructs 35S::HA:bZIP63:GFP (overexpression) and pbZIP63::HA:bZIP63:GFP (native promoter-driven expression). (B) Relative expression of *bZIP63* and its target genes *AT2G30600* and *BAM9* at EN in two *bzip63-2* lines carrying *35S::HA:bZIP63:GFP*, *bZIP63-ox3* and *bZIP63-ox4*. (C) Western blot analysis of HA:bZIP63:GFP accumulation in, *bZIP63-ox3* and *bZIP63-ox4,* at the EN. The fusion protein has ∼69 kDa. Total protein loaded was 40 µg for Ws, *bZIP63-ox3* and *bZIP63-ox4*. Actin (∼42 kDa), served as loading control and was used for normalization. Coomassie blue–stained gel shows Rubisco’s large subunit (RbcL, of ∼53 kDa). The ratios of the HA/ACTIN signals are indicated below the blots. (D) Relative expression of *bZIP63* and its target genes *AT2G30600* and *BAM9* at EN in *bzip63-2* lines carrying *pbZIP63::HA:bZIP63:GFP, bZIP63-co1*, *bZIP63-co2* and *bZIP63-co3*. (E) Western blot analysis of HA:bZIP63:GFP accumulation in *bZIP63-co1*, *bZIP63-co2* and *bZIP63-co3* at the EN. Total protein loaded was 40 µg for *bZIP63-co1*, *bZIP63-co2* and *bZIP63-co3.* (See C for details). The ratios of HA/ACTIN signals are indicated below the blots. (F) Relative *bZIP63* mRNA levels, and abundance of bZIP63 protein in Ws, *bZIP63-co1* and *bZIP63-ox3*. *bZIP63* transcript values are relative to Ws at the EN, and protein values represent abundance relative to *bZIP63-co1* at the EN. (G) Western blot analysis of HA:bZIP63:GFP accumulation in *bZIP63-co1* and *bZIP63-ox3* at the ED and EN. (See C for details). Total protein loaded was 60 µg for *bZIP63-co1,* and 5 µg for *bZIP63-ox3*. The average ratios and standard deviation of HA/ACTIN signals are indicated below the blots. (H, I) Western blot and analysis showing HA:bZIP63:GFP accumulation in *bZIP63-co1* and *bZIP63-ox3* at the EN in two independent experiments (See C for details). Total protein loaded was 60 µg for *bZIP63-co1*, and 5 µg for *bZIP63-ox3*. The average HA signals are indicated below the blots. All plants were grown for 21 days in short photoperiod (8 h light / 16 h dark). Plants in B, C, D and E were grown in 100 μmol m**^−2^** s**^−1^** of light intensity. Plants in F, G, H and I were grown in 80 μmol m**^−2^** s**^−1^** of light intensity. Values are means of two to three, and three to five replicates per genotype for protein and transcript data, respectively, and error bars indicate standard deviation. Statistical differences were assessed by Student’s t-test (p<0.05), where in B and D, ‘a’ and ’b’ denote expressions significantly different from Ws and *bzip63-2,* respectively, and in F, H and I, an asterisk (*) denotes significant difference.

**Figure S11.**
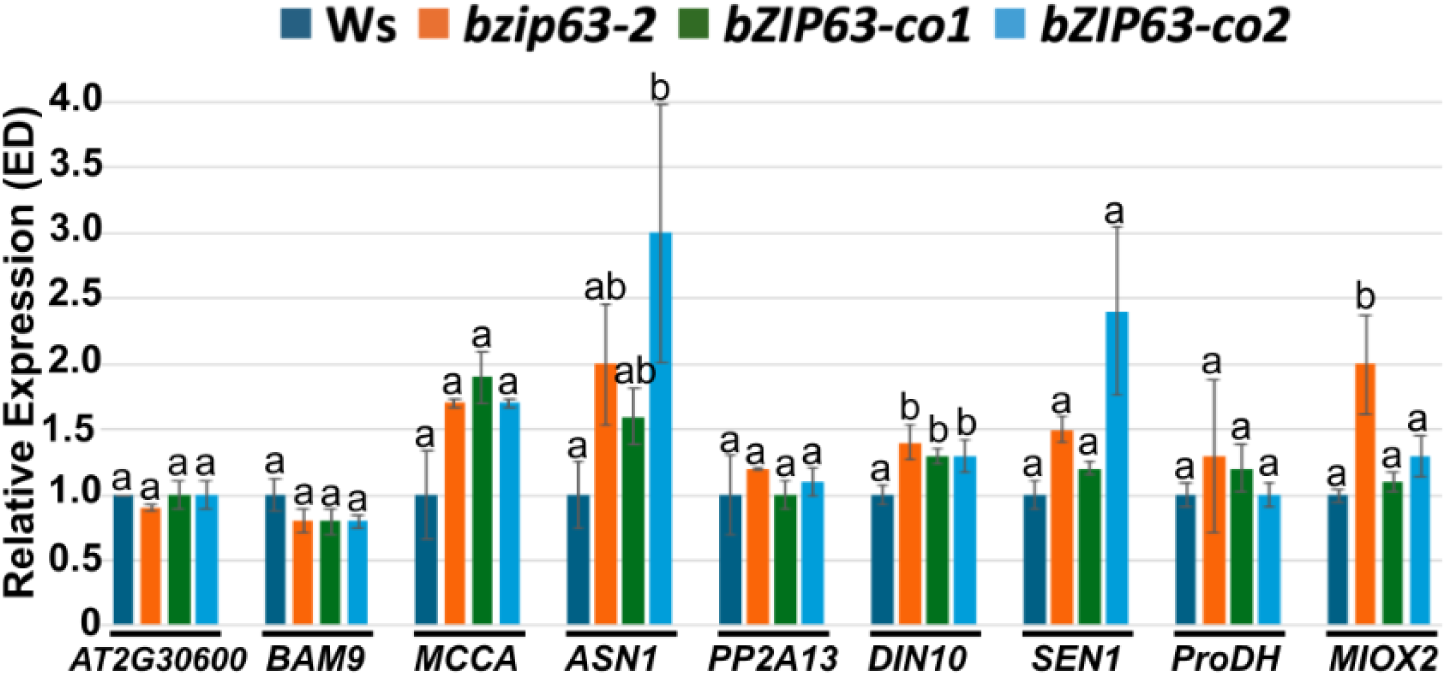
Relative expression of *bZIP63*-related SDIGs, *PP2-A13* and *BAM9* at ED in Ws, *bzip63-2*, *bZIP63-co1* and *bZIP63-co2*. Values represent expressions relative to the expression of each gene in Ws. Plants were grown for 21 days in short photoperiods in 80 μmol m^−2^ s^−1^ of light intensity. Values are means of three to five replicates per genotype, and error bars indicate standard deviation. Statistical differences were assessed by one-way ANOVA for parametric data, with different letters indicating significantly distinct groups (Tukey’s HSD test, p<0.05). For non-parametric data, the Kruskal-Wallis test was used, with different letters indicating significantly distinct groups (Dunn’s test, p < 0.05).

**Figure S12.**
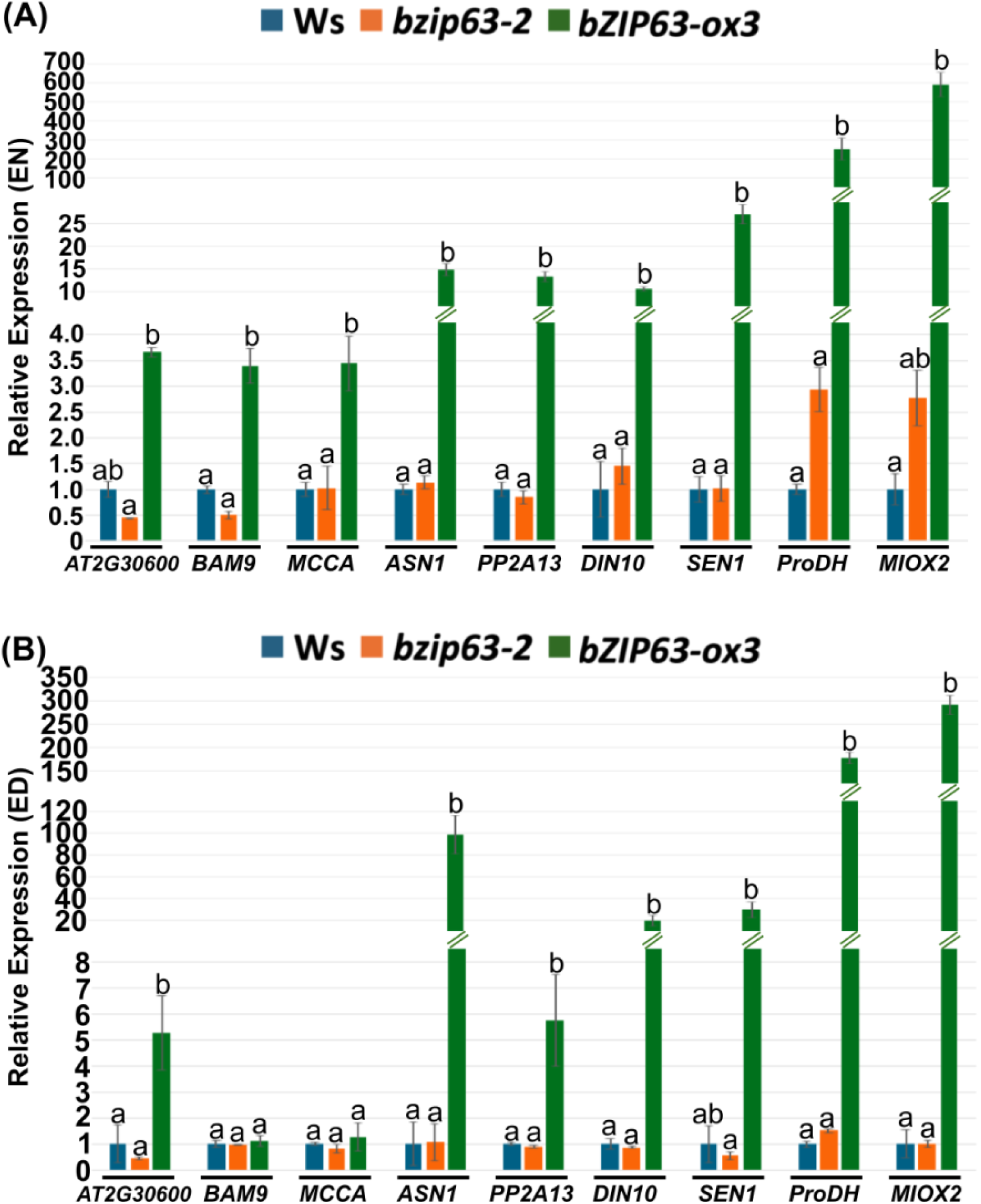
Relative expression of *bZIP63*-related SDIGs, *PP2-A13* and *BAM9* in Ws, *bzip63-2* and *bZIP63-ox3.* (A) at EN and (B) at ED. Values represent expressions relative to the expression of each gene in Ws and are means of three to five replicates per genotype. Error bars indicate standard deviation. Statistical differences were assessed by one-way ANOVA for parametric data, with different letters indicating significantly distinct groups (Tukey’s HSD test, p<0.05). For non-parametric data, the Kruskal-Wallis test was used, with different letters indicating significantly distinct groups (Dunn’s test, p<0.05). Plants were grown for 21 days in short photoperiods in 80 μmol m^−2^ s^−1^ of light intensity.

**Figure S13.**
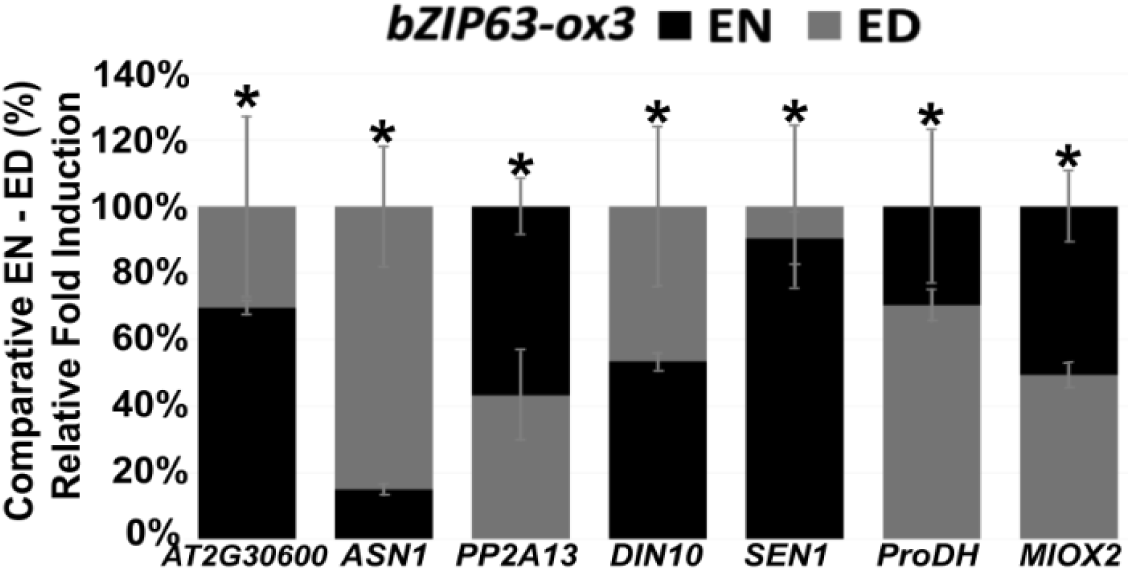
Fold induction of *bZIP63*-related SDIGs and *PP2-A13* in *bZIP63-ox3* relative to Ws at both EN and ED. The largest fold was set to 100% for each gene. Plants were grown for 21 days in short photoperiods and 80 μmol m^−2^ s^−1^ of light intensity. Values are means of three to five replicates, and error bars indicate standard deviation. Statistical differences were assessed by Student’s t-test (* p<0.05).

**Figure S14.**
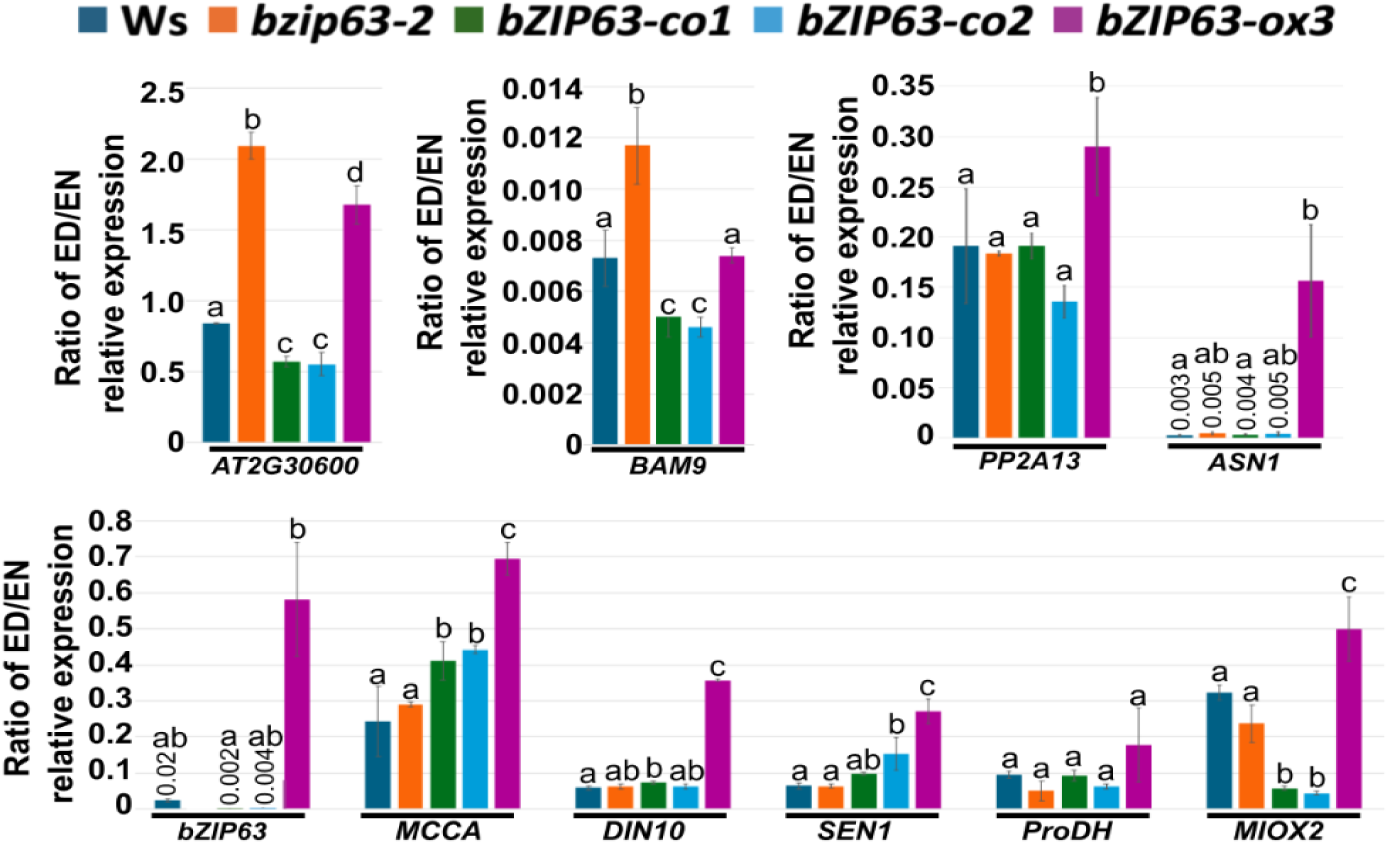
The EN–ED oscillatory expression pattern of *bZIP63*-related SDIGs, *PP2-A13*, and *BAM9* is preserved among Ws, *bzip63-2*, *bZIP63-co1*, *bZIP63-co2*, and *bZIP63-ox3*. Values represent the ratios of the average relative expression values of each gene in the different genotypes at ED relative to the average relative expression values at EN. Values are means, and error bars indicate standard deviation. Statistical differences were assessed by one-way ANOVA for ratios with parametric values, with different letters indicating significantly distinct groups (Tukey’s HSD test, p<0.05), and by Kruskal-Wallis, for ratios with non-parametric values, with different letters indicating significantly distinct groups (Dunn’s test, *p*<0.05). All plants were grown in short photoperiod (8 h light / 16 h dark), 80 μmol m**^−2^**s**^−1^** of light intensity for 21 days.

