## Supplementary Table S3 for "*bZIP63* misregulation affects growth and target gene expression under short-day photoperiods"

**Supplementary Table S3. Primers used in genotyping and gene construction**

| Primer | Sequence 5’-3’ | Purpose |
| --- | --- | --- |
| sgbZIP63_CISPR | GAATCGCAGTGCTTCCGAAT | Guide RNA used in the editing of the bZIP63 gene via CRISPR/Cas to obtain bzip63-5 and -15. |
| bzip63_CRISPR Fw | AATGCAAGACTGCCCAATCG | Identification of the bzip63-5 and bzip63-15 alleles. PCR is performed on the 5'UTR region of the gene, upstream of the indel. |
| bzip63_CRISPR Rv | GCTTGTTCTGATTCATTGGCTC | Identification of the bzip63-5 and bzip63-15 alleles. PCR is performed on the 2nd exon of the gene, downstream of the indel. |
| bZIP1-LP1/2 Fw | TCGTCATTCGATGAATCTTCC | Identification of the bzip1-1 allele. PCR is performed on the promoter region of the gene, upstream of the T-DNA insertion. |
| bZIP1-RP1 Rv | GCCATTTACATGCAAGGTACC | Identification of the bzip1-1 allele. PCR is performed on the promoter region of the gene, downstream of the T-DNA insertion. |
| bZIP53 Fw | TAAGGGAGAGCTAAGCCCATC | Identification of the bzip53-1 allele. PCR is performed on the promoter region of the gene, upstream of the T-DNA insertion. |
| bZIP53 Rv | TCGGATCATTATCGGATTCAG | Identification of the bzip53-1 allele. PCR is performed on the single exon region of the gene, downstream of the T-DNA insertion. |
| LBb1.3 | ATTTTGCCGATTTCGGAAC | Identification of T-DNA SALK mutants. PCR amplification at the left border of the T-DNA. |
| HygroFw#1 | AAGACCTGCCTGAAACCGAACT | Identification of the gene for hygromycin resistance (HygroR). |
| HygroRv#1 | AAGACCAATGCGGAGCATATAC | Identification of the gene for hygromycin resistance (HygroR). |
| bzip53_INRAE Fw | ACTCTTTCTCCGTCGTTTACCTC | Identification of the bZIP53 gene deletion. The primer is designed to anneal upstream of the coding region of the gene. |
| bzip53_INRAE Rv | AGAGGACTGAGGCTACAAAACC | Identification of the bZIP53 gene deletion. The primer is designed to anneal downstream of the coding region of the gene. |
| INRAE_bzip63 Fw | AAATGCAAGACTGCCCAATCG | Identification of the bZIP63 gene deletion. The primer is designed to anneal upstream of the coding region of the gene. |
| INRAE_bzip63 Rv | TCAACACAAACAATCAACGAAGA | Identification of the bZIP63 gene deletion. The primer is designed to anneal downstream of the coding region of the gene. |
| P_bZIP63_Fw1 | AAATGAGCTCTAAGCATCGTTCTCCCAAGAGA | Amplification of the bZIP63 promoter. |
| P_SpeI_bZIP63_Rv1 | AAAAACTAGTGTCTGATTATTACCCTAATGGCCC | Amplification of the bZIP63 promoter. |
