## Supplementary Table S4 for "*bZIP63* misregulation affects growth and target gene expression under short-day photoperiods"

**Supplementary table XXX.** Genes and primers used for gene expression quantification by RT-qPCR

| Gene | AGI | Primer sequence 5’-3’  Forward; Reverse | | Amplification efficiency  E =[ 10^(–1/slope)-1] |
| --- | --- | --- | --- | --- |
| AT1G13320 | AT1G13320 | CATGTTCCAAACTCTTACCTG  GTTCTCCACAACCGCTTGGT | 0.91 | |
| bZIP1 | AT5G4945 | TGTCGACGATCAGAACGCAA  AGGACGCCATTGGTTGTAGAG | 0.91 | |
| bZIP53 | AT3G62420 | GGCTTCGGAGTTGACGGATA  ATCTGCCAAGGGTTCTGCAT | 0.94 | |
| bZIP63 | AT5G28770 | GGAACTTTCATCAAACCTCAG  CACTGCTCATCATTGGTGTAG | 0.94 | |
| PRR7 | AT5G02810 | TTCCGAAAGAAGGTACGATAC  GCTATCCTCAATGTTTTTTATGT | 1.06 | |
| ASN1 | AT3G47340 | AGGTGCGGACGAGATCTTTG  GTGAAGAGCCTTGATCTTGC | 0.95 | |
| AT2G30600 | AT2G30600 | AAAGCCTATGCGGGTACTTC  CTGATGTTCTTCGCCTAAGTC | 1.04 | |
| BGAL4 | AT5G56870 | TTAACCTGGTACAAGTCTACG  GACGTCCAATATTTCTACCG | 0.93 | |
| ProDH | AT3G30775 | CGCTATACCGTATCTTCTCC  CTCTTAAGTTCCATCCTCATG | 1.1 | |
| BAM4 | AT5G55700 | TTACGTAGACAGATACATGATG  TCGGTTGCACACAGTTCTCT | 1.03 | |
| BAM9 | AT5G18670 | GAGCACCAATCACCTGAATC  CGACTCCTTGTTTCTTGCAG | 1.02 | |
| KMD4 | AT3G59940 | GGAAGGTAATTATGGATACGAT  CGTCGTCTTCACCATCATC | 0.95 | |
| DIN10 | AT5G20250 | TGTCAGTGATTCTCCTGGAA  CACCATCACGGGCAGGATC | 1.02 | |
| DIN4 | AT3G13450 | TCTTGACGCAGAAAACGAAGG  GCCCCAAAGCCTCCTGTAAC | 0.91 | |
| MCCA | AT1G03090 | TTAGGCCAAGCTGCTGTCTC  CTGTAGACGGGTGTTCATT | 0.99 | |
| BCAT1 | AT1G10060 | GGCTCTTCGTCGCTGCTTA  CATATTCTTCATCCTCACGTTCTGC | 1.0 | |
| TEM1 | AT1G25560 | CTGTTTTGAGAGCGCGTGAG  CTTCTCCGCGTGTTGTTTCG | 1.1 | |
| PP2-A13 | AT3G61060 | TGCTCGTCTCAACAGGATGT  ATTCCTGCTTTGTGCCATCGTC | 0.93 | |
| SEN1 | AT4G35770 | ACAGAGTCGGATCAGGAATGG  CTGTGATCGCCGTGAAGCC | 0.94 | |
| SEX1 | AT1G10760 | CCTCTACGACAGTGTACCAA  TCCAGCGCGTGCAATGTCT | 1.07 | |
| TPS9 | AT1G23870 | CTGAGCGACTGGCTATCTCC  ATCGTCTTCCATTCCGCCTC | 0.99 | |
| RVE7 | AT1G18330 | CGGCACATCCTTATCCTCGG  TTGGATGTCACTGGTACACGA | 0.91 | |
| PHYA | AT1G09570 | GAGCTGACTGGTCTTTCGGT  CATTCTGCTCCTCAGTTCCTTCT | 0.91 | |
| MIOX2 | AT2G19800 | CAACACCTTCTCCAAACCGC  ACTGGAAATGTATCGCCAACG | 0.98 | |
| BT2 | AT3G48360 | TGACGCCGAATCGAGGAAGA  ACTACGTTTGATGGACCGACC | 0.97 | |
